# Taming complexity in enzymatic saccharification: A predictive hierarchical modeling framework

**DOI:** 10.64898/2026.09.08.750015

**Authors:** Solmaz Hossein Khani, Ali Faraj, Grégoire Malandain, Gabriel Paës, Yassin Refahi

## Abstract

Lignocellulosic biomass biotechnological conversion relies on enzymatic hydrolysis of recalcitrant plant cell walls, which limits conversion efficiency and economic viability and renders sugar-release dynamics difficult to predict. Mathematical modeling has therefore emerged as a key approach for elucidating the mechanisms governing enzymatic hydrolysis and predicting its dynamics. Here, a hierarchical adsorption–inhibition framework is developed, beginning with a detailed Dynamic Adsorption–Inhibition Model (DyAIM). This model is then simplified into a Reduced Adsorption–Inhibition Model (ReAIM) by pruning weak adsorption and inhibition interactions. Finally, ReAIM is further simplified into an Effective Activity Model (EAM), in which nonproductive enzyme adsorption onto lignin is represented by a reduction in effective enzymatic activity. Plant cell wall composition is expressed as four structural polymers coupled to six soluble products, with inhibition by mono- and oligosaccharides acting on five functional enzyme pools. Using glucose, xylose and mannose release at two enzyme loadings, the model hierarchy is validated in reproducing saccharification dynamics and in delivering a quantitative product–enzyme inhibition network consistent with literature trends. The model framework was further validated by prediction of saccharification dynamics at an unseen enzyme loading, demonstrating robust performance beyond the calibration conditions. Sensitivity analysis further supported the progressive simplifications adopted in ReAIM and EAM. Maintaining predictive performance across increasing levels of simplification indicates that the hierarchy effectively distinguishes essential mechanisms from dispensable complexity, enabling rational model selection.

## 1 Introduction

Biotechnological conversion of lignocellulosic biomass relies critically on the enzymatic hydrolysis of plant cell walls [11]. Effective enzymatic hydrolysis is essential for achieving high yields and ensuring the economic viability of the process. Lignocellulosic plant biomass is composed mainly of cellulose, hemicelluloses, and lignin. Cellulose is a linear polysaccharide composed of D-glucose monomers linked by *β*(1 → 4) glycosidic bonds. Bundles of aligned *β*(1 → 4)-linked D-glucose chains form cellulose microfibrils, which contain both crystalline and amorphous regions. The crystalline region exhibits a high degree of structural order, contributing substantially to cellulose recalcitrance. The amorphous region is less ordered, exhibits weaker hydrogen bonding, and is more susceptible to hydrolytic deconstruction. Hemicelluloses represent an amorphous heterogeneous group of polysaccharides (e.g., xylans, mannans, galactans) composed of various sugar monomers, including pentoses (e.g., arabinose and xylose) and hexoses (e.g., mannose, and galactose) [44]. Lignin is a structurally complex aromatic biopolymer that provides hydrophobicity and mechanical strength to cell walls [67].

Lignocellulosic biomass is commonly classified as hardwoods, softwoods, or grasses. Hardwoods (e.g., poplar, oak) generally contain more cellulose and less lignin than softwoods, with xylan as the predominant hemicellulose. Softwoods (e.g., pine, spruce) are instead rich in galactoglucomannan as the main hemicellulose [29], whereas in grasses and grass-derived feedstocks (e.g., switchgrass, wheat straw), hemicelluloses are dominated by xylan [50].

Efficient enzymatic hydrolysis of lignocellulosic biomass into monosaccharides is challenging and requires the synergistic action of enzymes including cellulases (cellobiohydrolases, endoglucanases, and *β*-glucosidases) [9] together with hemicellulases. The latter include a set of backbone-cleaving enzymes, like xylanases, and mannanases, which target xylan, and mannan, respectively, and also debranching and deacetylating enzymes that hydrolyze side chains and esters [17].

Cellulose hydrolysis by cellulases proceeds through adsorption of enzymes onto the cellulose surface (often via carbohydrate-binding modules (CBMs)), formation of an enzyme–substrate complex, and catalytic cleavage of *β*-glucosidic bonds, where endoglucanases create new chain ends, exoglucanases release cellobiose processively, and *β*-glucosidases convert soluble oligosaccharides and cellobiose to glucose, followed by enzyme desorption back into the liquid phase [22]. In parallel, nonproductive binding to lignin, hemicelluloses, or nonproductive sites on cellulose, as well as product inhibition by mono- and oligosaccharides can significantly reduce overall hydrolysis efficiency [35]. In addition, hydrolysis efficiency is impeded by recalcitrance, arising from diverse features such as hierarchical wall architecture, cellulose crystallinity, and lignin–carbohydrate complexes (LCCs) that limit enzyme access to cellulose [40]. As a result, sugar release dynamics are difficult to predict *a priori*, even under well-controlled laboratory conditions, and remain challenging to extrapolate across enzyme loadings, substrates, and process conditions [21].

Mathematical modeling of enzymatic hydrolysis of lignocellulosic biomass has emerged as a valuable approach for elucidating the underlying mechanisms governing this complex process [8, 22]. Existing models can be divided into three categories: i) empirical models [42, 58]; ii) enzyme–substrate interaction models [4, 25]; iii) geometry- and substrate-structure-based models [32, 51]. Most of these models focus mainly on the enzymatic hydrolysis of cellulose. Empirical models rely on experimental data rather than mechanistic details. They include static correlations, linking hydrolysis yield at fixed times to substrate properties, and compact kinetic functions, describing yield over time without mechanistic details. Their strengths are simplicity and broad applicability with minimal data, but they lack mechanistic interpretability and perform poorly outside calibration. Thus, they are mostly useful for descriptive analysis, benchmarking, and techno-economic modeling. Enzyme–substrate interaction models describe how cellulases bind, act, and are hindered at insoluble interfaces by combining inhibition kinetics with adsorption thermodynamics and, in some cases, explicit processivity. Product inhibition is typically attributed to glucose and cellobiose acting on *β*-glucosidases, endoglucanases, and cellobiohydrolases. Enzyme binding is modeled using equilibrium isotherms, most commonly the Langmuir isotherm [69], or kinetic formulations [51]. Semi-mechanistic frameworks integrate these elements (surface reactions, Langmuir adsorption, and product inhibition) to fit reactor-scale sugar release [4, 25], whereas more mechanistic models add processivity, traffic effects, and transitions between productive and inactive bound states, thus accounting for early fast rates followed by slowdowns [36, 43]. Geometry- and substrate-structure-based models describe biomass hydrolysis by explicitly linking enzymatic activity to particle morphology and substrate architecture. By incorporating evolving structural features, such as cellulose degree of polymerization, porosity, and accessible surface area, these models capture how enzyme action is coupled to dynamic changes in biomass structure. Representative families of substrate-structure-based models include population balance models, which track the size distribution of cellulose fragments during hydrolysis [32, 41], and image-based and diffusion–reaction models, which describe enzymatic hydrolysis using imaged cell wall architecture [39, 51]. More recently, 4D (3D + time) imaging has extended such image-based approaches by characterizing, at the microscale, how cell wall architecture evolves during enzymatic hydrolysis [20, 47].

While mathematical modeling has been effective in translating the mechanisms of lignocellulosic enzymatic hydrolysis into a quantitative framework, most mechanistic models have focused primarily on cellulose hydrolysis and have treated hemicelluloses in a lumped manner. The composition of the enzyme cocktail is also commonly simplified. Similarly, product inhibition is often only partially considered, for example by accounting for the effects of cellobiose and glucose on cellulases while overlooking the broader network of inhibitory interactions and their relative strengths. Nonproductive enzyme adsorption to lignin is often represented by a global correction factor, without accounting for differences in affinity. Consequently, the coupled dynamics of enzyme adsorption, product inhibition, and substrate evolution remain only partially resolved and the level of mechanistic details required to achieve robust predictive power remains unclear.

This study presents a hierarchical modeling framework for enzymatic hydrolysis that integrates functional cellulolytic, xylanolytic, and mannanolytic enzyme pools, their adsorption onto cellulose, hemicelluloses, and lignin, their catalytic action in polysaccharide hydrolysis, and a detailed inhibition network accounting for the effects of mono- and oligosaccharides. The initial adsorption and inhibition structure is then systematically reduced by pruning weakly supported adsorption links and restricting the inhibition network to the strongest and most consistently observed interactions. The framework is validated using enzymatic hydrolysis data from sodium-chlorite–treated spruce wood.

## 2 Materials and methods

### 2.1 Determination of polymer-class composition from monosaccharide profiles

This study investigates the enzymatic hydrolysis of pretreated spruce wood, as raw spruce exhibits very limited yields [21]. The composition of the pretreated spruce wood was determined in our earlier work and was dominated by glucose (Glc, 58.89%), mannose (Man, 15.01%), lignin (11.30%), xylose (Xyl, 6.50%), galactose (Gal, 1.44%), arabinose (Ara, 0.90%), and glucuronic acid (GlcA, 0.35%), where the values in parentheses indicate the corresponding abbreviations and mass fractions [21]. Minor amounts of other sugars (e.g., fucose and rhamnose) and extractives were also present. These monosaccharides and lignin fractions were grouped into four polymer classes: cellulose, lignin, hemicelluloses whose hydrolysis mainly releases Man (assigned to O-acetyl-galactoglucomannan), denoted by GGM, and hemicelluloses whose hydrolysis mainly yields Xyl (assigned to arabino-4-*O*-methyl-glucuronoxylan), denoted by AGX [18].

In the subsequent mathematical model, cellulose, GGM and AGX serve as substrates for cellulolytic and hemicellulolytic enzymes. In assigning the monomeric sugars to polymer classes, Gal was mapped to GGM side chains, while GlcA and Ara were mapped to AGX non-backbone residues; Xyl and Man were considered as the respective backbone sugars. Glc originating from hemicellulose was attributed to GGM by adopting a representative softwood backbone ratio of Glc : Man = 1 : 4 [49], and the remaining Glc was assigned to cellulose. The minor fraction composed of minor sugars (e.g., fucose and rhamnose), and extractives were left unassigned. Using these rules, the polymer-class mass fractions were calculated as detailed in Section S1 in the supplementary material. Accordingly, the pretreated spruce contained 55.14% cellulose, 20.20% GGM, 7.65% AGX, and 11.30% lignin. These fractions sum to 94.29% of the total mass, with a residual fraction of 5.71%. For modeling purposes, polymer fractions were normalized to sum to 100%, resulting in 58.49% cellulose, 21.42% GGM, 8.11% AGX, and 11.98% lignin, denoted by 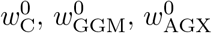, and 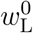, respectively.

### 2.2 Enzyme activity assays and conversion yields

Enzyme activities in the cocktail produced by *Trichoderma reesei* (provided by IFPEN, Rueil-Malmaison, France) were determined at 50 °C in 50mM citrate buffer (pH 4.8). Total cellulase activity was measured by the filter paper unit (FPU) assay [5]. Xylanase activity was determined using the Kidby assay, and endo-1,4-*β*-mannanase activity was quantified by a konjac glucomannan-based assay [26]. Activities of *β*-glucosidase, *β*-xylosidase, *α*-arabinosidase, and acetyl esterase were determined using pNP-glucopyranoside, pNP-xylopyranoside, pNP-arabinofuranoside, and pNP-acetate assays, respectively [46]. For reference, measured activities (mean ± standard deviation) were: cellulase 2.70 ± 0.47 FPU/mL; xylanase 9.33 ± 0.14 IU/mL; endo-1,4-*β*-mannanase 0.94 ± 0.14 IU/mL (konjac glucomannan); *β*-glucosidase 5.10±0.07 IU/mL; *β*-xylosidase 3.25±0.04 IU/mL; *α*-arabinosidase 1.60±0.04 IU/mL; and acetyl esterase 1.76 ± 0.12 IU/mL.

Sugar conversion yields at enzyme loadings of 15 and 30 FPU/g biomass were determined in our previous study [21]. Briefly, sample sections were placed in 200 µL mini-reactors, each containing 60 µL enzyme solution with enzyme loadings of 15 FPU/g biomass, or 30 FPU/g biomass, or acetate buffer for controls (i.e., datasets collected in the same conditions but without enzyme) and incubated at 50 °C; each condition was performed in triplicate. Hydrolysates were sampled at 0 h, 15 min, 1 h, 2 h, 4 h, 8 h, 12 h, 16 h, 20 h, and 24 h. For validation, for this study a separate experiment was conducted with an enzyme loading of 7.5 FPU/g biomass and hydrolysates were sampled at 0 h, 15 min, 1 h, 2 h, 4 h, and 8 h.

### 2.3 Model development: Dynamic Adsorption–Inhibition Model (DyAIM)

The modeling framework is constructed in three successive levels of detail. First, a detailed model is developed, in which enzyme adsorption/desorption on all relevant components and competitive product inhibition are represented explicitly. This detailed model is referred to as the *Dynamic Adsorption– Inhibition Model* (DyAIM). Its adsorption and inhibition networks were established through a literature survey; the selected interactions and the rationale for their inclusion are described in detail in the Results section (Sections 3.3.1 and 3.3.2, respectively). In a second step, the DyAIM structure is simplified by pruning weakly supported adsorption links and product inhibitions, generating the *Reduced Adsorption– Inhibition Model* (ReAIM, Section 2.4). Finally, a further simplified model variant, named *Effective Activity Model* (EAM, Section 2.5) is introduced, in which dynamic nonproductive enzyme adsorption to lignin is replaced by reduced effective enzyme activities. In this section, DyAIM is presented.

#### 2.3.1 Structural components and state variables

The hydrolyzed sample is represented by four components—cellulose, lignin, GGM, and AGX. Here, “GGM” and “AGX” denote the polymer classes, while *H*_GGM_ and *H*_AGX_ are the corresponding insoluble hemicellulose concentrations in the model. *C* and *L* represent the cellulose and lignin concentrations, respectively, as well as their corresponding polymer classes. Each component is modeled through an insoluble, time-dependent mass concentration, expressed on a volumetric basis (mg/mL) as *C*(*t*), *H*_GGM_(*t*), *H*_AGX_(*t*), and *L*(*t*). The total concentration of insoluble solids is then *S*(*t*) ≡ *C*(*t*) + *H*_GGM_(*t*) + *H*_AGX_(*t*) + *L*(*t*). For an initial biomass concentration *S*(0) (mg/mL), the component concentrations are 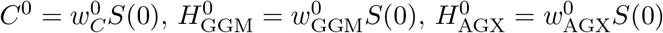, and 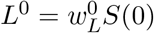.

During hydrolysis, lignin is assumed to remain chemically unchanged but may adsorb enzymes. In addition to the four insoluble components, the model explicitly tracks six soluble product pools in the liquid phase: cellobiose, with concentration G_2_; Glc, with concentration G_1_; xylooligosaccharides, denoted by XOS; mannan/glucomannan-derived oligosaccharides, denoted by MOS; Man, with concentration M_1_; and Xyl, with concentration X_1_. The symbols G_2_, XOS and MOS denote both the corresponding soluble product pools and their concentrations, with the intended meaning determined by the context. The MOS pool comprises soluble oligosaccharides originating from GGM, which are treated as a single mannan/glucomannan-derived oligosaccharides pool in the model. Each product is represented by a state variable with unit of mass concentration (mg/mL).

Only Glc, Xyl and Man were retained as explicit monosaccharide state variables. The low-abundance sugars (*<* 2.5 % of the dry mass) were assigned to side-chain residues of the hemicellulose polymers (GGM and AGX) and were therefore accounted for implicitly through the polymer-class mass balances. The model does, however, include the soluble intermediates that arise from depolymerization—G_2_ from cellulose, XOS from AGX and MOS from GGM—to capture the key steps of enzymatic hydrolysis.

#### 2.3.2 Enzyme pools

The enzyme cocktail is divided into five functional activity pools, E_1_–E_5_. Each pool is defined on an activity-concentration basis expressed as FPU/mL for cellulase activity and as IU/mL for all other activities.

- E_1_ (cellulolytic): exo- and endoacting cellulases that depolymerize cellulose, with soluble cellulose-derived products represented as G_2_.
- E_2_ (xylanolytic): endo-xylanases acting on the AGX backbone to produce XOS, together with associated debranching activities, including *α*-arabinosidase and acetyl esterase that facilitate AGX depolymerization.
- E_3_ (mannanolytic): composite mannanolytic activity, representing endo-1,4-*β*-mannanase together with exoacting mannanolytic/*β*-mannosidase activity acting on GGM. In the model, E_3_ catalyzes both the interfacial depolymerization of GGM into MOS and their subsequent conversion in the liquid phase, producing predominantly Man and also a smaller amount of Glc.
- E_4_ (*β*-glucosidase): soluble *β*-glucosidase converting G_2_ to Glc in the liquid phase.
- E_5_ (*β*-xylosidase): soluble *β*-xylosidase converting XOS to Xyl in the liquid phase.

The total activity concentration of each pool per slurry volume, denoted by *E*_*i*, tot_ (FPU/mL for *i* = 1, and IU/mL for *i* = 2, …, 5), is assumed to remain constant during enzymatic hydrolysis. Here, “slurry volume” refers to the total volume of the solid–liquid reaction mixture. Let *η* denote the enzyme cocktail dose (dimensionless volume-fraction expressed as mL of enzyme cocktail per mL of slurry), and let *A*_*i*_ (FPU/mL for *i* = 1 and IU/mL for *i* = 2, …, 5) denote the measured activity concentration of pool *i* (see Section S2 in the supplementary material for details about assignment of measured activities). Then, the total activity concentrations are expressed as *E*_*i*, tot_ = *η A*_*i*_, *i* = 1, …, 5. The enzyme cocktail dose *η*, expressed as a volume-fraction, was 0.0245, 0.049, and 0.098 for the datasets collected at 7.5, 15, and 30 FPU/g biomass, respectively.

#### 2.3.3 Adsorption kinetics

Enzyme adsorption is described by dynamic Langmuir kinetics [31], with adsorbed enzyme expressed as activity equivalent. Thus, the adsorbed enzyme variable represents bound catalytic activity per unit mass of solid, rather than enzyme mass per unit mass of solid. For each enzyme pool *i*, E_*i*_, 1 ≤ *i* ≤ 5, the set of possible binding components is denoted by 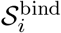 with 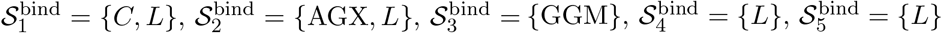.

Let *E*_*i,f*_ (*t*), *i* ∈ {1, 2, 3, 4, 5}, denote the liquid-phase free activity concentration of E_*i*_ at time *t*, expressed in FPU/mL for *i* = 1 and IU/mL for *i* = 2, …, 5. Let 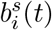 denote the adsorbed activity equivalent of enzyme pool *i* on *s*, expressed per unit mass of the corresponding component (FPU/mg for *i* = 1 and IU/mg for *i* = 2, …, 5). 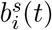 evolves according to

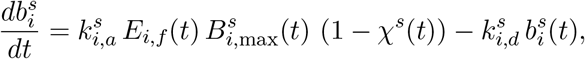

where 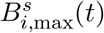 is the maximum activity-equivalent adsorption capacity for E_*i*_ on *s*, expressed per unit mass of the corresponding component, with the same unit as 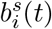. The parameters 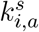 and 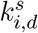 are the adsorption and desorption rate coefficients, respectively, with 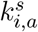 expressed in mL/(FPU h) for *i* = 1 and in mL/(IU h) for *i* = 2, …, 5, and 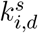 expressed in h^*−*1^. The term *χ*^*s*^(*t*) denotes the fractional occupancy of *s*. Let 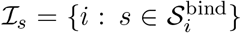 be the set of enzyme pools that may bind to *s*. The fractional occupancy is then

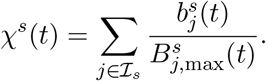

For lignin, the nonproductively adsorbing enzyme pools are assumed to compete for a shared pool of lignin adsorption sites. Thus, *χ*^*L*^(*t*) denotes the total fractional occupancy of the shared lignin surface with 0 ≤ *χ*^*L*^(*t*) ≤ 1, rather than a collection of independent pool-specific occupancy, and is given by

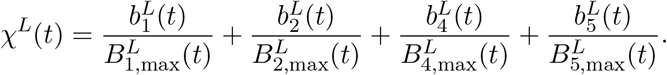

This shared-occupancy formulation prevents the lignin surface from being saturated independently by each enzyme pool and ensures that 1 − *χ*^*L*^(*t*) represents the remaining vacant fraction of the common lignin binding capacity. The free liquid-phase activity of each enzyme pool is determined by the corresponding activity concentration balance

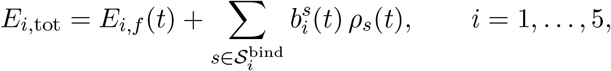

where *ρ*_*s*_(*t*) is the concentration of the solid component *s*. At *t* = 0, all enzymes are initially unbound, i.e., 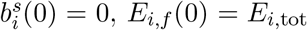. In the activity concentration balances, the product 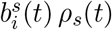 represents the volumetric bound activity of enzyme pool *i* associated with *s*.

The adsorbed-activity variables are therefore 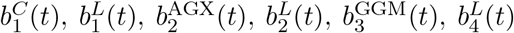, and 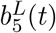. See Section S3 in the supplementary material for the explicit equations.

#### 2.3.4 Coupling adsorption capacities to substrate evolution

In DyAIM, the maximum adsorption capacities of cellulose and lignin are assumed to evolve with the extent of hemicelluloses depletion to account for changes in substrate induced by hemicelluloses removal. The time-dependent maximum adsorption capacities are given by

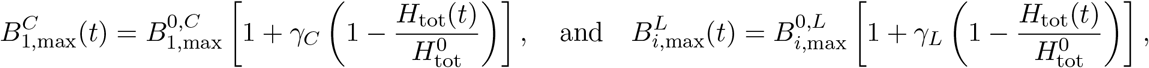

for *i* ∈ {1, 2, 4, 5}. *H*_tot_(*t*) = *H*_AGX_(*t*)+*H*_GGM_(*t*), and 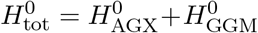 denote the total hemicellulose concentration and its initial value, respectively. The dimensionless parameters *γ*_*C*_ ≥ 0 and *γ*_*L*_ ≥ 0 control the increase in adsorption capacity of cellulose and lignin, respectively, as hemicelluloses are depleted. 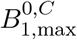, and 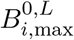, *i* ∈ {1, 2, 4, 5}, denote the initial maximum adsorption capacities.

The maximum adsorption capacities of AGX and GGM are assumed constant, i.e., 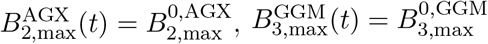, where 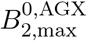, and, 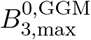 denote the baseline adsorption capacities at *t* = 0. The linear dependence on hemicelluloses depletion was chosen as the simplest nontrivial formulation that preserves the initial capacities, ensures positive capacities for *γ*_*C*_, *γ*_*L*_ ≥ 0, and limits the number of additional parameters. For all enzyme pool *i* and component *s*, 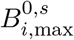 has the same unit as 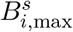.

#### 2.3.5 Reaction rates and product inhibition

Hydrolysis reaction rates are described separately for heterogeneous and homogeneous pathways, with product inhibitions incorporated as competitive inhibitions.

Heterogeneous hydrolysis rates are assumed proportional to the productively bound enzyme activity and to the corresponding polymer-class mass concentration. Competitive product inhibition is included in the denominator. Below are these heterogeneous reactions.

- Cellulose hydrolysis by E_1_ is inhibited competitively by G_2_, Glc, Xyl, Man, XOS, and MOS:

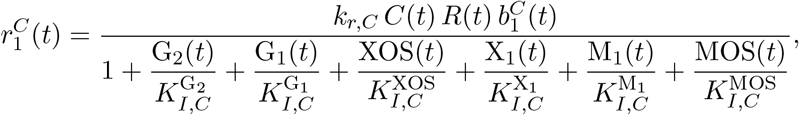

where 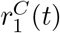 (mg/(mL h)) is the reaction rate, *k*_*r,C*_ (mg/(FPU h)) is the effective hydrolysis rate coefficient and 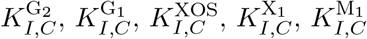, and 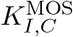 are the competitive-inhibition constants for G_2_, Glc, XOS, Xyl, Man, and MOS, respectively. The inhibition constants have the same units as their corresponding product concentrations (mg/mL). The dimensionless factor *R*(*t*) = (*C*(*t*)*/C*^0^) ^*α*^ represents the reactivity factor with the dimensionless exponent *α* and describes how cellulose reactivity scales with its hydrolysis.
- Hydrolysis of AGX by E_2_ is inhibited competitively by XOS, G_2_, Glc, and MOS:

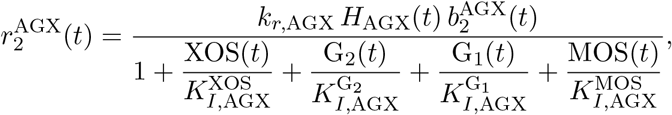

where 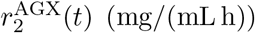 is the reaction rate, 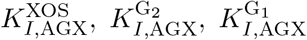, and 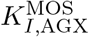 are the competitive inhibition constants (expressed in mg/mL) for XOS, G_2_, Glc, and MOS, respectively, and *k*_*r*,AGX_ is the effective hydrolysis rate coefficient.
- Hydrolysis of GGM by the mannanolytic pool E_3_ is inhibited by Man:

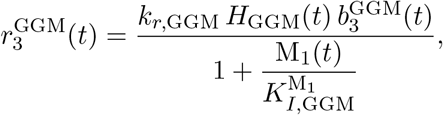

where 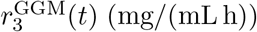 is the reaction rate, *k*_*r*,GGM_ is the effective hydrolysis rate coefficient and 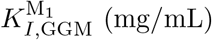 is the inhibition constant for Man. Homogeneous reactions in the liquid phase are represented by Michaelis–Menten kinetics with competitive inhibition.
- G_2_ hydrolysis to Glc by E_4_ is inhibited by Glc and G_2_:

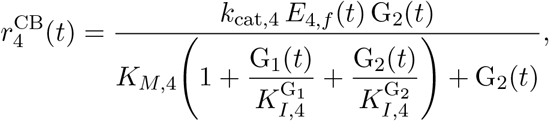

where 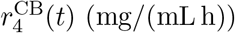 is the reaction rate, *k*_cat,4_ is the catalytic constant, *K*_*M*,4_ is the Michaelis constant, and 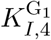and 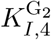 are the competitive inhibition constants for Glc and G_2_, respectively. The Michaelis and inhibition constants are expressed in units of mg/mL.
- Hydrolysis of XOS to Xyl by E_5_ is inhibited by Xyl:

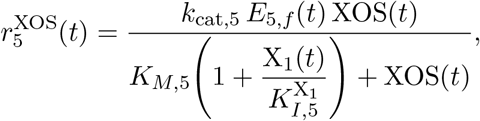

where 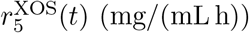 is the reaction rate, 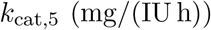 is the catalytic constant, *K*_*M*,5_ (mg/mL) is the Michaelis constant, and 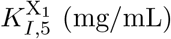 is the Xyl inhibition constant.
- Hydrolysis of MOS to Man by the mannanolytic pool E_3_ is inhibited by Man:

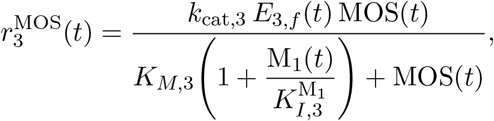

where 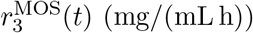 is the reaction rate, *k*_cat,3_ is the catalytic constant, *K*_*M*,3_ (mg/mL) is the Michaelis constant, and 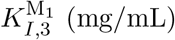 is the Man inhibition constant.

#### 2.3.6 Component mass balance equations

The mass balance equations listed below follow from the principle of mass conservation. The coefficients incorporate both hydration associated with conversion of anhydro residues to hydrated soluble products and the restriction of tracked soluble products to the explicitly modeled backbone-derived sugar pools.

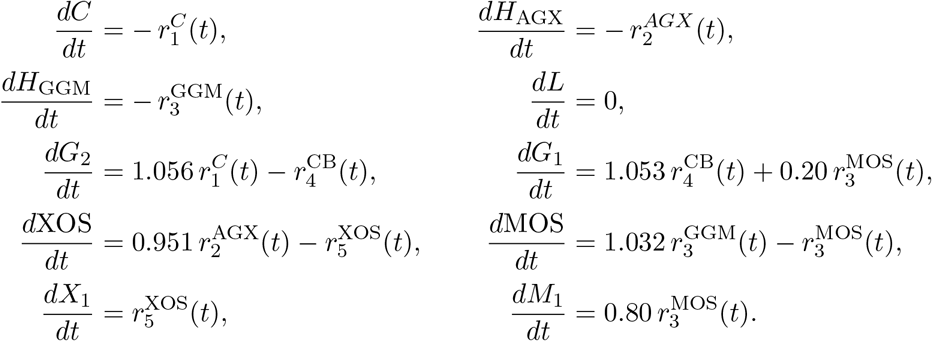

### 2.4 Model simplification: Reduced Adsorption–Inhibition Model (ReAIM)

DyAIM is simplified to the Reduced Adsorption–Inhibition Model (ReAIM) which retains only the strongest, best-supported enzyme adsorptions and the strongest product inhibitors for each enzyme pool. In addition, ReAIM assumes time-invariant adsorption capacities by fixing all maximum adsorption capacities to their baselines (i.e., 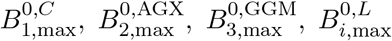 (*i* = 1, 4, 5)). All other components like state variables, and stoichiometric coefficients remain unchanged.

Specifically, ReAIM omits the nonproductive binding of E_2_ to lignin. Therefore the admissible binding sets are 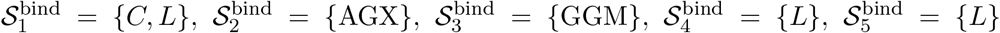. Dynamic adsorption follows the same Langmuir kinetics as in DyAIM with a simplified fractional occupancy for lignin

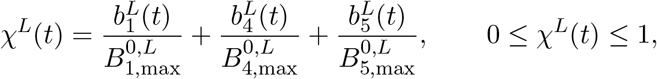

with the corresponding adsorption equations identical to those in DyAIM. As in DyAIM, the free liquid-phase activity concentration of each enzyme pool is obtained using the ReAIM admissible binding sets. See Section S4 in the supplementary material for the explicit adsorption and enzyme activity concentration balance equations. In addition, ReAIM retains only the dominant competitive inhibitors: E_1_ is inhibited by G_2_, XOS and MOS; E_2_ by XOS, G_2_ and MOS; E_3_ by Man; E_4_ by Glc; and E_5_ by Xyl. Thus, the heterogeneous reaction rates in ReAIM are

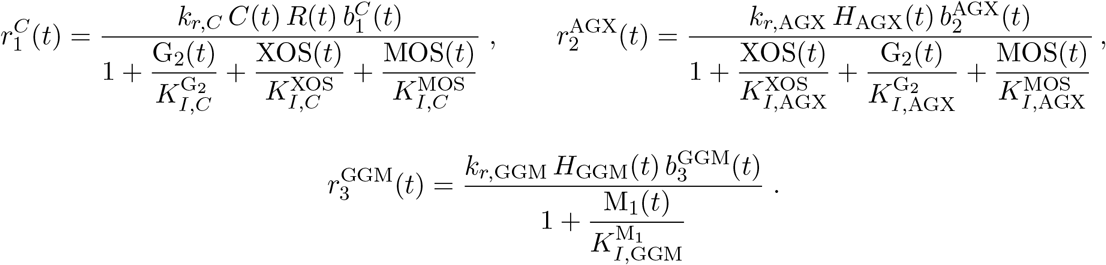

The homogeneous reactions rates for the solution-phase steps are

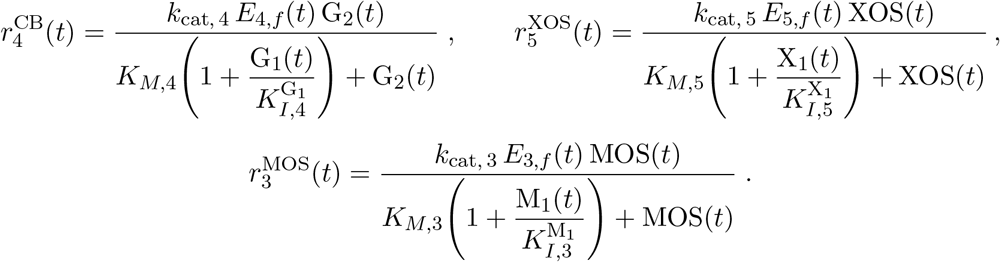

Here, *R*(*t*) and the parameter units are as defined in DyAIM. The component mass balances follow the DyAIM structure, using the simplified rate expressions defined above.

### 2.5 Effective Activity Model (EAM)

ReAIM is simplified into Effective Activity Model (EAM) which treats nonproductive lignin adsorption as a static reduction in catalytic activity concentration rather than a dynamic process. In EAM, explicit lignin-adsorption dynamics are replaced by an unavailable fraction, *ϕ*_*i*_, of the initial activity concentration of lignin-binding enzyme pool *i*. This reduction in activity concentration is assumed to occur instantaneously at the onset of hydrolysis and remain constant throughout the reaction. The effective catalytic activity concentrations are defined as *E*_*i*, eff_ = (1 − *ϕ*_*i*_) *E*_*i*, tot_, for *i* ∈ {1, 4, 5}, with 0 ≤ *ϕ*_*i*_ ≤ 1. No lignin-adsorption correction is applied to E_2_ and E_3_ (i.e., *ϕ*_2_ = *ϕ*_3_ = 0), thus *E*_*i*, eff_ = *E*_*i*, tot_, *i* ∈ {2, 3}. Accordingly, *E*_*i*,eff_ replaces *E*_*i*,tot_ in the enzyme mass balances used to determine the free enzyme activities, while dynamic adsorption is restricted to productive binding. The heterogeneous and homogeneous reaction-rate expressions and component mass balances retain the same functional form as in ReAIM. Explicit equations for productive adsorption and free enzymes are provided in Section S5 in the supplementary material.

### 2.6 Parameter estimation

A warm-start parameter-estimation strategy, using estimates from simpler variants to initialize the parameter estimation of more complex ones, was used to exploit the hierarchical relationship among the three model variants. The models were calibrated sequentially, beginning with EAM. The resulting EAM parameter estimates were used to initialize ReAIM calibration, whose optimized parameters were subsequently used to initialize DyAIM calibration (Figure 1.A). Thus, parameters shared across model variants inherited their initial values from the preceding model. Within each model variant, the optimization process involved a sequential, staged approach to parameter estimation (Figure 1.B). The first three stages each focused on parameters associated with the yield of a specific monosaccharide, in the following order: Man, Glc, and Xyl. During these stages, the residuals were weighted differently to preserve the fit quality of sugars optimized in earlier stages. In the final stage, all remaining parameters were optimized simultaneously, with equal weights assigned to the residuals of each sugar to obtain a comprehensive fit. In addition, the parameters were constrained to be identical for the two enzyme loadings of 15 and 30 FPU/g biomass and only enzyme cocktail dose, (i.e., *η*) changed. Calibration was performed against experimentally measured Glc, Xyl, and Man conversion yields, expressed as percentages of the theoretical maximum. For methodological details of multi-stage procedure see Section S6 and Table S1 in the supplementary material.

**Figure 1:**
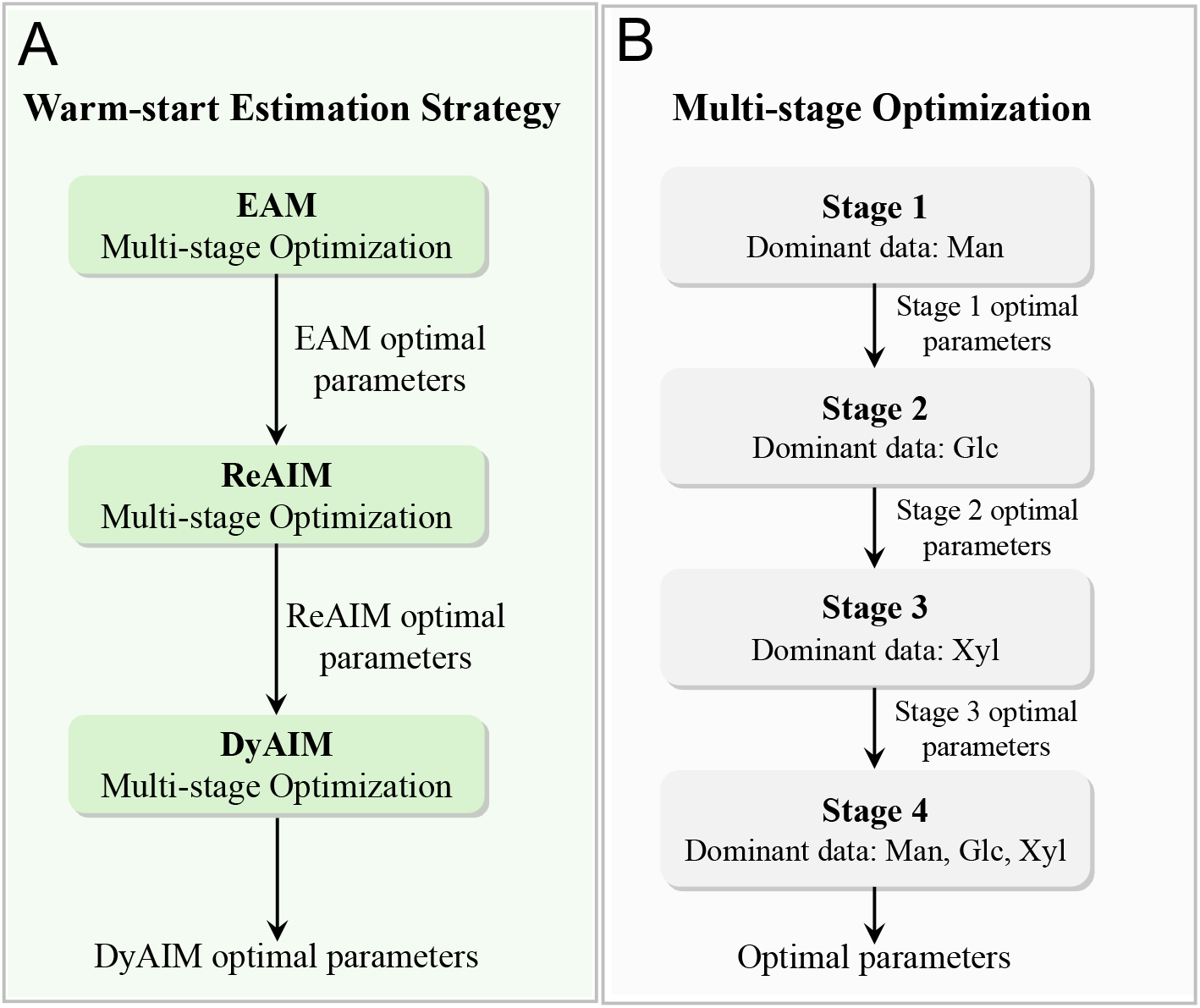
Warm-start parameter estimation with staged-procedure workflow. (A) The warm-start estimation, in which optimized parameters are transferred sequentially from EAM to ReAIM and then to DyAIM. (B) Within each model variant, parameters are estimated using a multi-stage procedure. At each stage, only a subset of parameters is estimated, and the resulting estimate are transferred to the subsequent stages.

## 3 Results and discussion

### 3.1 Inferring patterns of enzyme–lignin adsorption

Nonproductive enzyme adsorption onto lignin is a major sink for catalytic capacity during hydrolysis [67]. The literature was therefore surveyed to determine, for each enzyme class, whether adsorption onto lignin has been reported to be sufficiently strong and consistently reported. The reported magnitude of enzyme adsorption onto lignin varies substantially among studies. This variability reflects differences in lignin characteristics, including its abundance, spatial distribution, composition, and physicochemical features like molecular weight, which determine lignin affinity for nonproductive adsorption [57, 67]. Likewise, the type and properties of enzymes such as the presence or absence of carbohydrate-binding modules, strongly influence the extent of lignin–enzyme interactions [16]. Furthermore, cellulase–lignin interactions are further influenced by factors such as enzyme source, processing conditions (e.g., pH and temperature), and the severity of pretreatment [57, 62]. Consequently, the available evidence was interpreted qualitatively to identify enzyme–lignin interactions that could be considered non-negligible.

Post-hydrolysis analyses often have reported that 40–50% (in some cases, an even greater fraction) of the cellulases associated with cellobiohydrolases and endoglucanases can remain adsorbed to the lignin-rich residue [24]. Substantially lower adsorption levels have nevertheless been reported in other studies (typically *<* 15%) [33]. Despite this quantitative variability, adsorption of both cellobiohydrolases and endoglucanases onto lignin has been repeatedly observed. The available evidence therefore supports the presence of non-negligible lignin adsorption for both cellulase classes.

Significant variability has been reported for *β*-glucosidases. Earlier work reported that *β*-glucosidases from Novozym 188 adsorb only weakly to lignin, whereas *β*-glucosidases in more recent commercial preparations such as Cellic CTec2 exhibit strong lignin adsorption [24]. High lignin affinity has similarly been reported for some *β*-glucosidases [33]. By contrast, other studies have indicated that *β*-glucosidase activity is less strongly affected by lignin than the activities of core cellulases [16]. Thus, although the extent of adsorption varies with the origin and properties of *β*-glucosidase, the available evidence suggests that lignin adsorption is generally non-negligible.

Xylanases also exhibit substantial variability in adsorption behavior: whereas certain isoforms bind strongly to lignin, others display little to no affinity [16, 24, 33]. Notably, studies examining the adsorption of *Trichoderma reesei* enzymes on softwood lignin report very low levels of xylanase adsorption [33]. Because the xylanase considered in the present enzyme cocktail is produced by *Trichoderma reesei*, and because the model is applied to a softwood-derived substrate, this enzyme–lignin interaction was considered weak under the conditions represented in the model.

Most adsorption studies have concentrated on cellobiohydrolases, and endoglucanases, *β*-glucosidases and xylanases. As a result, the adsorption behavior of other functionally important enzymes, such as mannanases and *β*-xylosidases, remains comparatively under-characterized, and quantitative adsorption data for accessory hemicellulases are still limited. Within the limited evidence available, mannanase appears to exhibit only minimal adsorption [33], whereas *β*-xylosidase has been reported to adsorb strongly to lignin [61].

Taken together, the literature survey indicates that, within an enzymatic cocktail composed of cellobiohydrolases, endoglucanases, *β*-glucosidases, *β*-xylosidases, endo-xylanases, mannanases, and *β*-mannosidases, most components exhibit the potential to bind to lignin to varying extents. Notable exceptions are mannanases, for which adsorption appears negligible, and xylanases (produced by *Trichoderma reesei* and investigated on softwood lignin), for which only low levels of lignin binding have been reported.

### 3.2 Inferring product inhibition network

In addition to nonproductive adsorption of enzymes on lignin, activity of the enzymes is also limited by inhibition from the reaction products. The available literature was surveyed to identify the principal product inhibitors of each enzyme class in the cocktail and to assess their relative inhibitory strengths. Overall, monomeric sugars (such as Glc, and Xyl), disaccharides (such as G_2_) and oligomeric sugars (such as xylooligomers) inhibit enzymes during lignocellulosic biomass enzymatic hydrolysis, but the degree of inhibition varies significantly between types of sugars. Oligomeric sugars generally inhibit more strongly than the monomeric sugars which can exceed monomer inhibition by over 100-fold [28], with the notable exceptions of *β*-glucosidases and *β*-xylosidases, for which Glc and Xyl, respectively, are typically the predominant product inhibitors [1, 57]. To examine these overall trends at the enzyme level, the available evidence was analyzed separately for each enzyme class. The resulting inhibition patterns are detailed below and reported in Table 1.

**Table 1:** Summary of product inhibition during lignocellulosic biomass enzymatic hydrolysis.

| Enzyme class | Principal product inhibitors | Inhibition mode | Relative inhibitory strength (qualitative) | Notes with citations |
| --- | --- | --- | --- | --- |
| Cellobiohydrolases (CBHs) | XOS; G<sub>2</sub>; manno-oligosaccharides; Glc; Xyl; Man | Competitive (XOS, G<sub>2</sub>, manno-oligosaccharides; Glc for CBHs) | XOS strongest; G<sub>2</sub> strong; Glc weaker than G<sub>2</sub>; Xyl < XOS/G<sub>2</sub>/Glc; Man < Xyl | XOS often shows higher affinity than G<sub>2</sub> [28, 45]; G<sub>2</sub> is a canonical strong inhibitor [38]; Glc competitively inhibits however weaker than G<sub>2</sub> [2, 7]; Xyl and Man exhibit weaker inhibitions [23, 45]; manno-oligosaccharides also cause significant competitive inhibition [64]. |
| Endoglucanases (EGs) | G<sub>2</sub>; XOS; Glc (indirect via β-glucosidase inhibition) | Competitive (G<sub>2</sub>, XOS) | G<sub>2</sub> strong; XOS strong; Glc weak directly | G<sub>2</sub> inhibits via active site/CBM [6, 12, 13, 30]; Glc indirectly increases G<sub>2</sub> by inhibiting β-glucosidases [3]; XOS competitive and strong [15]; monomeric Xyl/Man do not directly inhibit major cellulolytic enzymes but can reduce productive binding/processivity [54, 68]. |
| Endo-xylanases | G<sub>2</sub>; Glc; XOS; manno-oligosaccharides; Xyl (study-dependent) | Competitive (G<sub>2</sub>, Glc, XOS) | G<sub>2</sub> > Glc > cellulose; XOS strong and increases with DP; manno-oligosaccharides strong (smaller DP, stronger inhibition) | Xyl: inhibition/no effect/stimulation reported [10, 59, 65]; XOS competitive, potency increases with DP; generally stronger than cello-oligosaccharides [37]; monomeric Man does not inhibit [68]. |
| β-glucosidases (BGs) | Glc; Xyl (variable); Man (conflicting); G<sub>2</sub> (less than EGs/CBHs); xylan/xylooligomers/mannan (no effect in some studies) | Competitive for Glc; Xyl competitive at high concentration in some BGs | Glc strong; Xyl variable; Man variable and generally ≤ Glc | Preferentially inhibited by Glc [2]; G<sub>2</sub> more strongly affects EGs/CBHs [13]; many reports show no inhibition by Xyl/xylooligomers/xylan/Man/Gal [45, 63, 64]; Xyl can stimulate at low-moderate levels and inhibit competitively at high levels in some BGs [55, 56]; Man: conflicting reports (strong vs. no effect) [52, 68]. |
| β-xylosidases | Xyl; (Glc sensitivity for GH3 family only) | Generally competitive (Xyl) | Xyl strong; Glc/G<sub>2</sub> usually none (except GH3) | Xyl competitive inhibitor [60, 66]; Glc/G<sub>2</sub> generally no inhibition, except GH3 enzymes [34]; no evidence for inhibition by manno-oligosaccharides; other monosaccharides show no effect [27]. |
| Endo-mannanases | Man | Competitive | Man moderate-strong; others not reported | Man is the only demonstrated inhibitor among putative products; no direct evidence for inhibition by Glc, Xyl, XOS, manno-oligosaccharides, or G<sub>2</sub> [14]. |

- Among inhibitors of cellobiohydrolases (CBHs), XOS exhibits the strongest inhibition, binding competitively with affinities often higher than those of G_2_ [28, 45], the canonical strong inhibitor [38]. Glc is also recognized as an inhibitor [2]. However, G_2_ is a markedly stronger inhibitor of cellulase activity than Glc, although both act as competitive inhibitors of CBHs [7]. Xyl also inhibits CBHs, but its effect is generally weaker than that of xylooligomers, G_2_, and Glc [45]. Similarly, Man inhibits CBHs, but to a lesser extent than Xyl [23]. Manno-oligosaccharides inhibit CBHs as well, acting as competitive inhibitors and significantly reducing cellulase activity [64].
- Endoglucanases (EGs) activity is strongly inhibited by G_2_ through competitive inhibition and only weakly by Glc [6, 12, 30]. Cellobiose inhibits EGs via binding to the active site and/or the carbohydrate-binding module [13]. Glc has a significant indirect effect on EGs by inhibiting *β*-glucosidase, and causing G_2_ to accumulate, which then strongly inhibits EGs [3]. EGs are strongly inhibited by XOS through competitive binding [15]. Hemicellulose-derived monosaccharides have also been reported to affect cellulase activity [54]; however, monomeric sugars such as Xyl and Man do not appear to directly inhibit the major cellulolytic enzymes, including EGs, *β*-glucosidases, xylanases, and CBHs in cellulase mixtures [68]. Nonetheless, these monomers can reduce the overall efficiency of cellulose hydrolysis by reducing the productive binding and processivity of cellulases [68].
- The hydrolysis products G_2_, and Glc negatively affect the hydrolytic activity of endo-xylanase through competitive inhibition, whereas *β*-xylosidase remains largely unaffected. Among these, G_2_ has the strongest inhibitory effect on endo-xylanase, followed by Glc [10]. The effect of Xyl is inconsistent across studies: some report strong inhibition [10], others find no inhibition [65], and some even observe stimulation of enzyme activity [59]. XOS act as competitive inhibitors of endo-xylanase, with inhibitory strength increasing with degree of polymerization (DP), and are generally more potent inhibitors than cello-oligosaccharides [37]. Manno-oligosaccharides can also act as strong inhibitors of xylanase activity with manno-oligosaccharides with fewer subunits causing stronger inhibition of xylanases compared to the larger manno-oligosaccharides [15] while it has been reported that monomeric Man does not inhibit xylanase activity [68].
- *β*-glucosidase activity is preferentially inhibited by Glc [2], whereas G_2_ more strongly affects exo- and endoglucanases [13]. Xyl does not universally inhibit *β*-glucosidase activity. Absence of inhibitory effect of xylooligomers, and Xyl, Man, and Gal on *β*-glucosidase has been reported [45, 63, 64]. At low to moderate concentrations, Xyl can stimulate activity in some *β*-glucosidases [56], while at high concentrations, it may act as a competitive inhibitor [55]. Inhibition of *β*-glucosidase by manno-oligosaccharides has not been reported; however, *β*-glucosidase activity was found to be unaffected by mannan [64]. There is clear conflicting findings regarding Man’s inhibitory effect on *β*-glucosidase activity, with some studies reporting strong inhibition [52] and others reporting no effect [68]. This suggests that the response to Man might depend on the specific *β*-glucosidase source. The degree of inhibition is also generally less than that observed with Glc.
- Xyl is generally a competitive inhibitor of *β*-xylosidase [60, 66]. Glc and G_2_ generally do not inhibit *β*-xylosidase activity [10] with the exception of *β*-xylosidases of GH3 family which is sensitive to Glc [34]. There is no direct evidence or data supporting inhibition of *β*-xylosidase by manno-oligosaccharides. Furthermore, beyond Xyl, the monosaccharides Man, Ara, Gal, Glc, and fructose have shown no effect on *β*-xylosidase activity [27].
- Among putative reaction products, Man is the only one with demonstrated inhibitory effects on endo-mannanase, acting as a competitive inhibitor [14]. Man also acts typically as a competitive inhibitor of *β*-mannosidases, though the extent of inhibition varies considerably across enzyme sources and families [53]. By contrast, Glc, Xyl, XOS, manno-oligosaccharides, and G_2_ have no direct experimental evidence supporting inhibition of endo-mannanase activity to the best of our knowledge.

### 3.3 Hierarchical modeling framework for enzymatic hydrolysis

Building on the adsorption and inhibition interactions outlined above, three nested dynamic models of spruce saccharification were developed: the Dynamic Adsorption–Inhibition Model (DyAIM), the Reduced Adsorption–Inhibition Model (ReAIM), and the Effective Activity Model (EAM). DyAIM was a full reference model in which all plausible adsorption links and product inhibitors were included. ReAIM and EAM were then derived through successive structural simplification while retaining the dominant physicochemical mechanisms. Specifically, ReAIM pruned weak adsorption and inhibition links and EAM replaced dynamic enzyme–lignin binding with fixed reductions in effective enzyme activity.

#### 3.3.1 Adsorption network in the model hierarchy

In DyAIM, adsorption was represented explicitly as a bipartite network between the five functional enzyme pools (E_1_–E_5_) and cellulose, arabino-4-*O*-methyl-glucuronoxylan (AGX), galactoglucomannan (GGM), and lignin, with enzyme–lignin interactions represented in line with inferred adsorption patterns by including all non-negligible interactions (Figure 2.A.1). Accordingly, the cellulolytic pool E_1_ was allowed to bind productively to cellulose and nonproductively to lignin; the xylanolytic pool E_2_ to bind productively to AGX and nonproductively to lignin; the mannanolytic pool E_3_ to bind productively to GGM only; and the E_4_ and E_5_ pools (*β*-glucosidase and *β*-xylosidase) to bind nonproductively to lignin only. Dynamic Langmuir kinetics were used to describe productive and nonproductive adsorptions.

**Figure 2:**
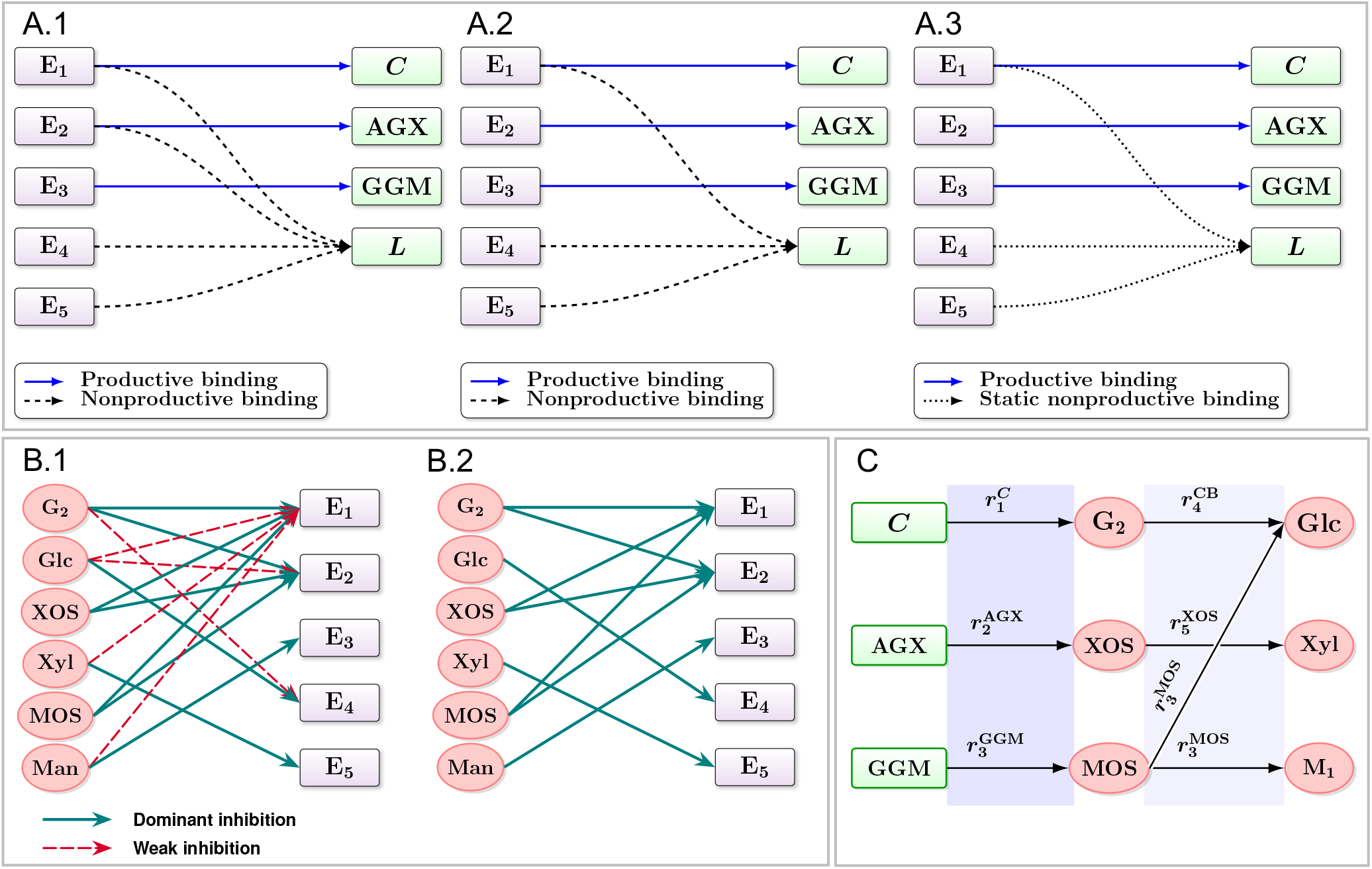
(A) Schematic representation of adsorption networks across the model hierarchy. (A.1) DyAIM adsorption network (A.2) ReAIM adsorption network. (A.3) EAM adsorption network. Purple and green rectangles represent enzyme pools and solid-phase components, respectively; blue solid, black dashed, and black dotted arrows indicate productive, dynamic nonproductive, and static nonproductive adsorptions, respectively. (B) Product–inhibition networks. (B.1) Product–inhibition network in DyAIM (B.2) Product–inhibition network in ReAIM and EAM. Red ellipses denote products. Green solid arrows and red dashed arrows represent dominant and weak inhibitions, respectively. (C) Representation of heterogeneous and homogeneous hydrolysis reactions. Heterogeneous reactions are shown in the left shaded panels. Homogeneous reactions are shown in the right shaded panels.

Productive polysaccharide sites were considered as enzyme-pool specific, such that no competition between pools for cellulose, AGX, or GGM was introduced, whereas all pools adsorbing to lignin competed for a shared pool of lignin adsorption sites. Adsorption capacities and substrate evolution were explicitly coupled; as hemicelluloses were solubilized, the adsorption capacities of cellulose and lignin were allowed to increase. DyAIM thus provided a deliberately rich adsorption network with evolving exposure of cellulose and lignin in which all plausible enzyme–component interactions were included and allowed to compete dynamically through shared lignin adsorption sites.

ReAIM was derived from DyAIM by pruning weak adsorption interactions. For E_2_, nonproductive lignin binding was removed so E_2_ could bind only productively to AGX (Figure 2.A.2). In addition, the model decouples lignin and cellulose adsorption capacities from hydrolysis of hemicelluloses. EAM collapses all lignin-bound enzyme states into fixed, enzyme-specific catalytically unavailable fractions; only productive polysaccharide adsorption stays dynamic (Figure 2.A.3).

#### 3.3.2 Product inhibition network in the model hierarchy

In DyAIM, a deliberately inclusive inhibition network was adopted according to the inferred inhibition network. Whenever the literature provided clear or plausible evidence that a given product inhibited a given enzyme pool, the corresponding inhibition interaction was included in the inhibition network (Figure 2.B.1). In this way, the model was constructed to represent the full range of reported inhibition patterns, including oligomer- and monomer-level effects on both cellulolytic and hemicellulolytic activities. Within this network, E_1_ was inhibited by G_2_, Glc, XOS, Xyl, MOS, and Man. G_2_, XOS, and MOS were considered as the dominant inhibitors, in line with reports indicating that oligomeric sugars inhibit cellulases more strongly than their monomeric counterparts [15, 28, 45], while the monomeric sugars Glc, Xyl, and Man were retained as weaker inhibitors to account for plausible but modest monomer effects. For E_2_, strong inhibition by XOS, G_2_, and MOS was included, together with weaker inhibition by Glc. By contrast, the mannanolytic pool E_3_ was taken to be inhibited only by Man, reflecting the consistent identification of Man as the principal competitive inhibitor of endo-mannanases and *β*-mannosidases. E_4_ was inhibited by Glc and weakly by G_2_, and E_5_ was inhibited by Xyl, while other potential inhibitors were not included due to limited or conflicting evidence.

ReAIM simplifies the DyAIM inhibition network by discarding the weak product-inhibition links (Figure 2.B.2). For E_1_, only the three oligomeric products G_2_, XOS, and MOS were kept as explicit inhibitors, whereas the weaker and less consistently reported effects of Glc, Xyl, and Man on cellulases were removed. For E_2_, a similar reduction was performed: inhibition by XOS, G_2_, and MOS was kept, while inhibition by Glc was not taken into account. Man remained the only inhibitor of E_3_. Glc was kept as the predominant inhibitor of E_4_ and Xyl continued to inhibit E_5_. EAM inherited this same reduced inhibition graph without further modifications; EAM differs only in the way enzyme-lignin adsorption is modeled, as previously explained.

Together, DyAIM, ReAIM, and EAM form a hierarchy with decreasing adsorption and inhibition network density. DyAIM uses a dense inhibition network where all plausible oligomers and monomers can inhibit associated enzyme pools. ReAIM focuses only on key inhibitors, and EAM keeps this compact structure while further simplifying nonproductive lignin binding.

#### 3.3.3 Reaction structure and full model specification

Beyond adsorption and inhibition, the three model variants share a common reaction skeleton. Reactions are divided into heterogeneous and homogeneous steps (Figure 2.C). Heterogeneous reactions describe the depolymerization of cellulose, GGM, and AGX by the enzyme pools E_1_–E_3_, with rates proportional to the corresponding solid concentration and productively bound activity, and modulated by competitive product inhibition. Homogeneous reactions describe the conversion of intermediate products in solution: G_2_ to Glc by E_4_, XOS to Xyl by E_5_, and MOS to Man (and, in fixed proportion, to Glc) by E_3_. These liquid-phase steps are written in Michaelis–Menten form with competitive inhibition. The functional forms of all heterogeneous and homogeneous rates are identical across DyAIM, ReAIM, and EAM. In addition, all models use the same reactivity factor, allowing cellulose reactivity to decrease with conversion. In DyAIM, the heterogeneous reaction rates depend on bound activities through a fully dynamic adsorption network combined with a dense product-inhibition network. In ReAIM, the same reaction expressions are driven by bound activities obtained from a reduced adsorption network and by a pruned inhibition network of dominant inhibitors. In EAM, the initial activities of the lignin binding enzyme pools are reduced by fixed lignin-adsorption fractions.

### 3.4 Saccharification dynamics are captured accurately by all three model variants

To estimate model parameters, a warm-start strategy was adopted which exploits the hierarchy–first fitting EAM, then using its optimized parameters to initialize ReAIM calibration, and finally using the ReAIM estimates to initialize DyAIM calibration. Fitting was systematically performed against the Glc, Xyl, and Man conversion yield dynamics measured at enzyme loadings of 15 and 30 FPU/g biomass. This simultaneous parameter estimation using both 15 and 30 FPU/ g biomass datasets aimed to reduce the risk of overfitting to a single condition and to provide a more rigorous assessment of whether models can capture the dependence of saccharification on enzyme loading. See Table S2 and Table S3 in the supplementary material for the experimentally determined and optimal parameter values, respectively. Technical details of the fit-quality assessment are provided in Section S7 in the supplementary material.

All three models accurately captured the saccharification dynamics (Figure 3 and Table S4 in the supplementary material). Across the pooled dataset combining both enzyme loadings, average *R*^2^ values were approximately 0.99. When evaluated separately, average *R*^2^ exceeded 0.98 at 15 FPU/g biomass and 0.99 at 30 FPU/g biomass, and remained high for individual sugars. Because these uniformly high *R*^2^ values provided limited discrimination, performance was compared primarily using range-normalized root mean square error (NRMSE) defined as RMSE divided by the observed range of the corresponding sugar–enzyme-loading block and expressed as a percentage.

**Figure 3:**
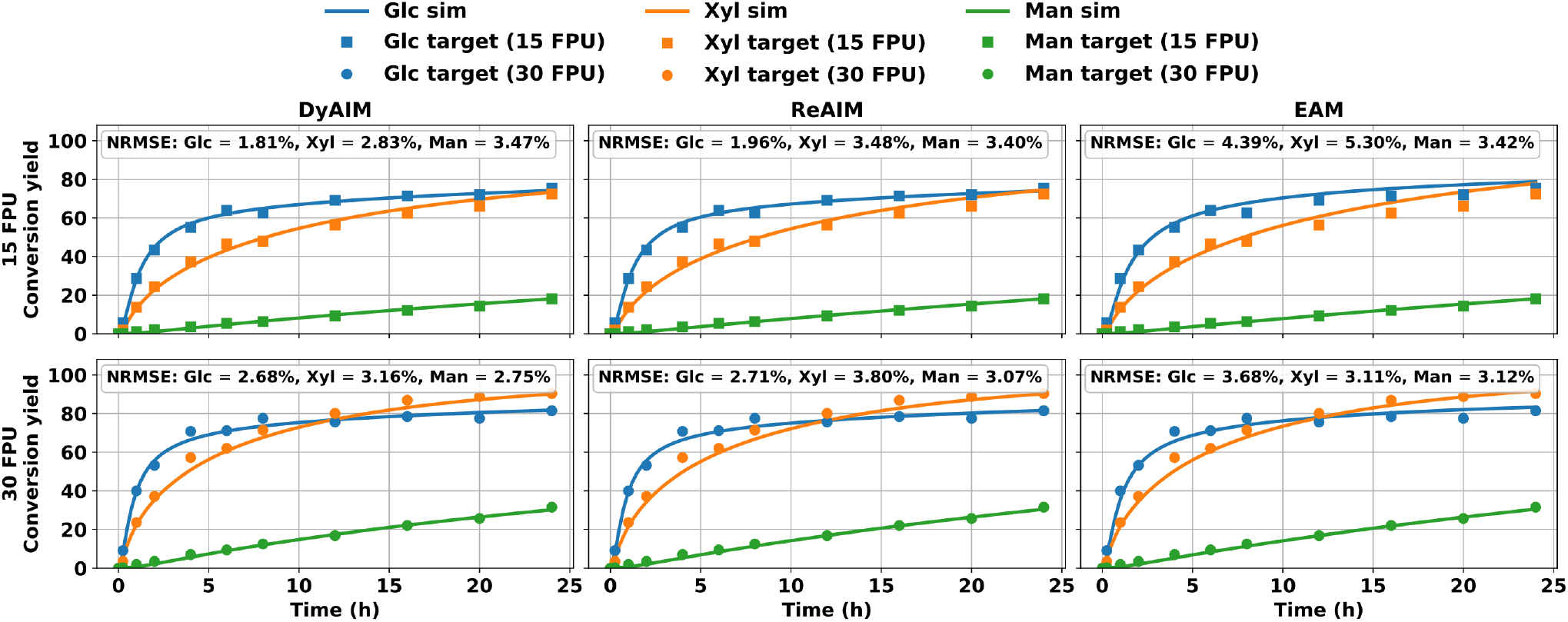
Model predictions versus experimental saccharification dynamics. Solid lines denote the simulated conversions of Glc (blue), Xyl (orange), and Man (green), while symbols denote the corresponding experimental data at enzyme loadings of 15 FPU/g biomass (squares) and 30 FPU/g biomass (circles), expressed as % of the maximum theoretical conversion. Columns correspond to the DyAIM, ReAIM, and EAM, with the upper and lower rows showing results for 15 and 30 FPU/g biomass, respectively.

The models yielded global NRMSE values (for all saccharification measurements obtained at both enzyme loadings) of 2.83% for DyAIM, 3.13% for ReAIM, and 3.93% for EAM. When evaluated separately at each enzyme loading, DyAIM yielded NRMSE values of 2.79% at 15 FPU/ g biomass and 2.87% at 30 FPU /g biomass, whereas ReAIM yielded corresponding values of 3.03% and 3.23%. EAM showed a higher error at the lower enzyme loading, with an NRMSE of 4.44% at 15 FPU/g biomass. However, its performance improved at 30 FPU/g biomass, where the NRMSE decreased to 3.31%, approaching the performance of DyAIM and ReAIM. Analysis of individual sugars showed a consistent ranking in model performance: DyAIM had the lowest NRMSE values, followed by ReAIM and then EAM. This ranking was consistent with the global NRMSE values, with EAM still showing robust performance. Importantly, this performance was obtained using a single set of parameters fitted simultaneously to both the 15 and 30 FPU/g biomass datasets, rather than fitting the model separately for each enzyme loading. This result indicates that the models capture the enzyme-dose dependence of enzymatic hydrolysis yield. Simultaneous calibration also facilitates the mechanistic interpretability of the fitted parameters, since differences between the two datasets can be attributed to enzyme loading rather than to parameter adjustments.

### 3.5 External validation at an unseen enzyme loading

To evaluate the models outside the calibration conditions, an external validation was performed at an enzyme loading of 7.5 FPU/g biomass. This dataset was not used in parameter estimation which relied only on the 15 and 30 FPU/g biomass datasets. At this enzyme loading, measurements were collected over the initial phase of the reaction, corresponding to the period of fastest saccharification in the Glc and Xyl release dynamics. This early phase is particularly informative for characterizing hydrolysis kinetics, because it captures the highest observed hydrolysis rates, enzyme–substrate interaction effects, and the rapid depletion of the most accessible sites. It is therefore arguably the most critical part of the trajectory from both a mechanistic and an industrial perspective. For each model, saccharification at 7.5 FPU/g biomass was simulated by modifying only the enzyme-cocktail dose, represented by the parameter *η*, while keeping the remaining parameter values inferred from the 15 and 30 FPU/g biomass datasets. Model predictions were subsequently compared with the experimentally measured Glc, Xyl, and Man conversion dynamics.

Overall, all three models showed good predictive ability (Figure 4). At the individual sugar level, differences in prediction error were observed. Across all models, Glc was predicted with the lowest error, with NRMSE values of 5.26%, 4.76%, and 2.35% for DyAIM, ReAIM, and EAM, respectively. Xyl showed intermediate errors, with NRMSE values of 6.13%, 5.41% and 7.37% for DyAIM, ReAIM, and EAM, respectively. In contrast, Man exhibited the largest errors, with NRMSE of 13.46%, 14.92% and 15.04% for DyAIM, ReAIM, and EAM, respectively. The larger errors for Man, however, should be interpreted in light of the low Man conversion at 7.5 FPU/g biomass, which was only 4.07%. At low conversion, the Man range is narrow, so even small absolute errors can result in relatively high NRMSE values. This interpretation is supported by the small absolute RMSE values for Man, ranging from approximately 0.55 to 0.61 acorss different model variants. By comparison, Glc and Xyl exhibited larger absolute RMSE values, ranging from 1.40 to 3.15 and from 1.67 to 2.28, respectively, while yielding lower range-normalized errors because their observed ranges were larger. Thus, the higher normalized error for Man mainly reflects its low conversion and narrow dynamic range, rather than a large absolute prediction error. The *R*^2^ values ranged approximately between 0.98 and 0.996 for Glc, between 0.96 and 0.98 for Xyl, and between 0.83 and 0.86 for Man.

**Figure 4:**
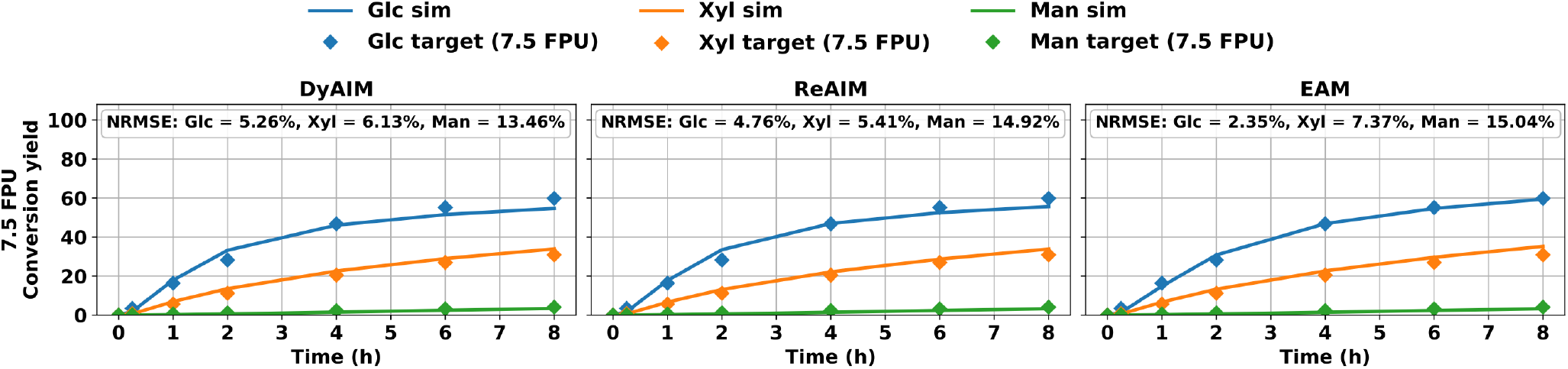
Model predictions versus unseen experimental saccharification data at an enzyme loading of 7.5 FPU/g biomass. Solid lines denote the predicted conversions of Glc (blue), Xyl (orange), and Man (green), while diamond symbols denote the corresponding experimental data, expressed as % of the maximum theoretical conversion.

When considering sugars collectively, the global NRMSE values were 9.07%, 9.57%, and 9.76% for DyAIM, ReAIM, and EAM, respectively. However, these global NRMSE values should be interpreted in relation to the individual sugar level results, since the relatively high global metric is partly driven by the elevated range-normalized errors for Man. When considering the predictive performance of individual model variants, EAM demonstrated highest accuracy in reproducing Glc dynamics with remarkably low NRMSE values of 2.35%. However, EAM exhibited larger errors for Xyl, with NRMSE values of 7.37%, and its performance for Man was comparable to that of the other models. These results demonstrate EAM’s ability to accurately predict Glc dynamics, supporting its robustness in capturing dynamics of Glc as the predominant sugar.

### 3.6 Model-inferred product inhibition strengths agree with literature trends

Following parameter estimation, the fitted inhibition constants were analyzed to infer the relative strengths of product inhibition. For each product–enzyme pair, inhibition strength was interpreted as inversely proportional to the corresponding inhibition constant, such that lower values indicated stronger inhibition (Table S5 in the supplementary material).

For E_1_, XOS emerged as the dominant inhibitor, with the ranking XOS *>* G_2_ *>* MOS ≫ Glc *>* Xyl ≈ Man. This inferred inhibition pattern was broadly consistent with the literature-based expectations (Table 1): E_1_ was most strongly inhibited by oligomeric products and G_2_, whereas the monomeric sugars exhibited comparatively weak inhibitions. In the case of E_2_, the relative strengths also point to dominant inhibition by hemicellulose-derived oligosaccharides where XOS was the strongest inhibitor, followed by MOS, with rankings XOS *>* MOS *>* G_2_ *>* Glc. The inhibition of E_3_ is governed by Man with relative strength depending on the reaction phase. Man exerts a stronger inhibitory effect on the heterogeneous GGM hydrolysis than it does on the homogeneous MOS hydrolysis reaction.

Glc exerts the strongest inhibition on E_4_, although its inhibitory effect is only slightly greater than that of G_2_.

For E_5_, Xyl is the only inhibitor considered in the models, precluding a comparison of relative inhibition strengths. Its accumulation may however be a key constraint on XOS conversion.

Overall, the fitted inhibition constants reflect the key qualitative features of the previously identified inhibition network. The predicted product inhibition strengths are consistent with trends reported in the literature: oligosaccharides are the dominant inhibitors of upstream heterogeneous deconstruction steps, whereas monosaccharides dominate inhibition of downstream reactions involving *β*-glucosidases and *β*-xylosidases, for which Glc and Xyl, respectively, are typically the principal product inhibitors.

### 3.7 Sensitivity analysis validates the staged estimation framework and model simplifications

A sensitivity analysis was conducted to examine the extent to which the model predictions are influenced by variations in the estimated parameters and to identify the parameters exerting the greatest influence on the predicted concentrations of Man, Glc, and Xyl. The corresponding sensitivity functions, which describe the local response of the predicted outputs to changes in individual model parameters (how a small change in a parameter affects the model predictions at a given time) are shown in Figure 5 (see Section S9 in the supplementary material for their mathematical definition). All sensitivity functions presented here were evaluated at the identified parameter vector. Changes in enzyme loading were found not to alter the *qualitative* behavior of the sensitivity functions, and only moderate effects were observed on their *quantitative* values. Similarly, when a parameter was shared by the DyAIM and a simplified model variant (ReAIM or EAM), the same qualitative influence of that parameter on a given model output was observed for both models (the DyAIM and the simplified variant). For these reasons, sensitivity functions are presented only for DyAIM at an enzyme loading of 15 FPU/g biomass.

**Figure 5:**
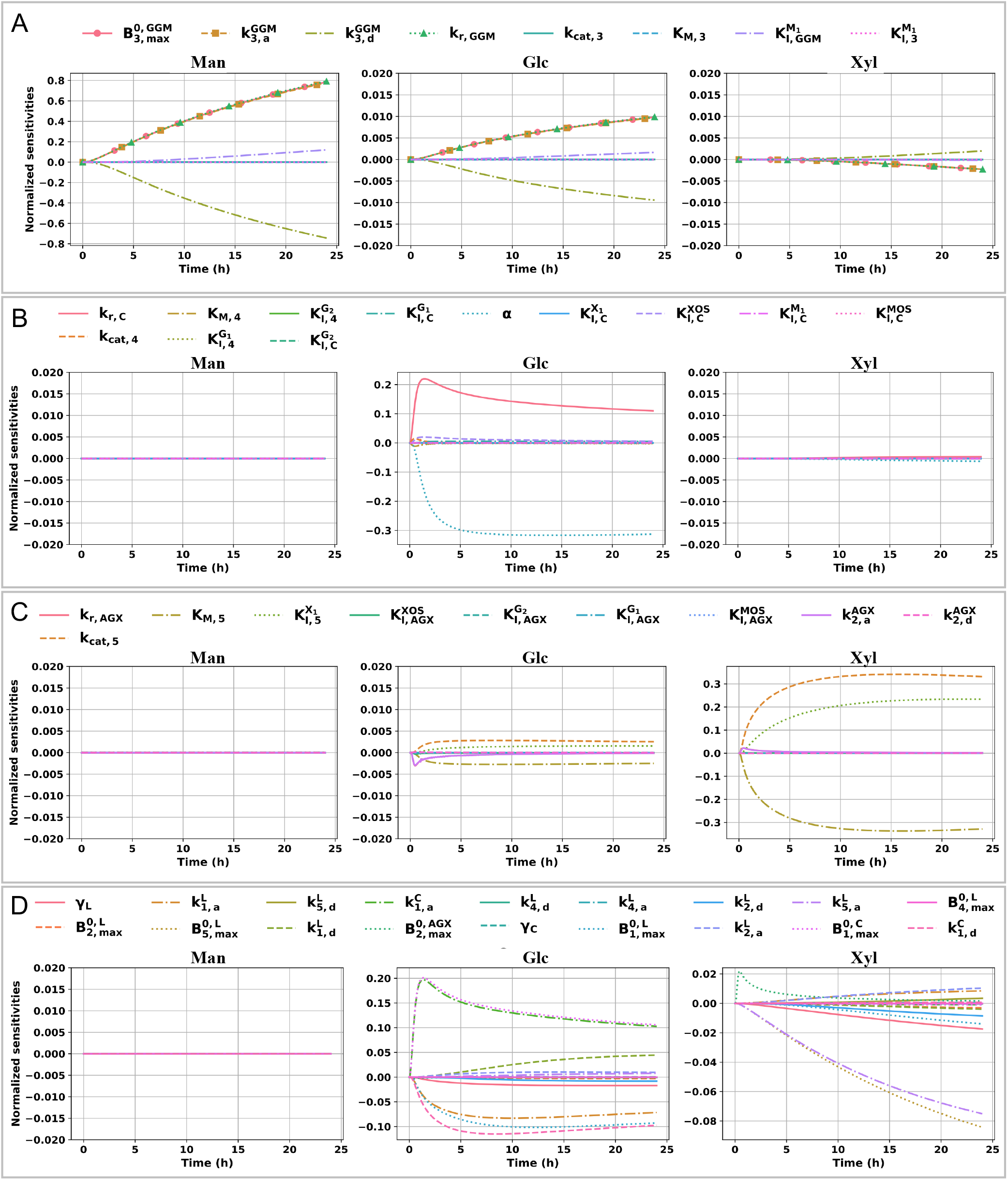
Time evolution of the sensitivity functions for DyAIM with enzyme loading of 15 FPU/g biomass. The sensitivity functions are grouped according to the subsets of free parameters estimated at each stage of the sequential multi-stage fitting procedure: (A), (B), (C), and (D) correspond to stages 1, 2, 3, and 4, respectively.

The sensitivity profiles indicate that Man conversion dynamics are primarily controlled by parameters associated with enzyme binding to GGM and the subsequent hydrolysis of GGM by the mannanolytic enzyme pool. More specifically, the parameters that have the strongest influence on Man conversion dynamics are 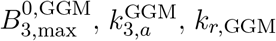, and 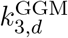. As shown in Figure 5.A, the corresponding Man sensitivity functions are almost linearly dependent, indicating that the effects of these parameters on the model output are difficult to distinguish independently. Consistent with this behavior, the estimated values of 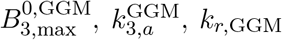, and 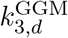 are strongly correlated. This correlation arises from compensatory effects among the parameters, where a change in one parameter can be offset by a change in another while yielding similar Man conversion dynamics. For example, an increase in 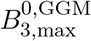 can be compensated by a decrease in 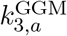, as can be inferred from the differential equation governing 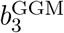. The parameter 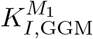 also influences Man conversion dynamics, although its effect is comparatively moderate.

The predicted Glc conversion dynamics are mainly determined by two parameters among those in the second stage of the multi-stage estimation approach with direct mechanistic relevance to cellulose conversion: the cellulose reactivity exponent and the cellulose hydrolysis rate coefficient. More specifically, the parameters *α* and *k*_*r,C*_ exert the strongest influence on the predicted Glc conversion dynamics, with *α* showing the largest sensitivity. The effects of the parameter *k*_*r,C*_ are particularly pronounced during the early phase of hydrolysis. This indicates that measurements collected during this initial period are especially informative for identifying this parameter. Accordingly, the denser sampling adopted during the first 8 hours in the present work is important for preserving the information content of the experimental data (Figure 5.B). The Xyl conversion dynamics is mainly influenced by *k*_cat,5_, *K*_*M*,5_ and 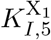, by order of significance, among the parameters estimated in the third stage. The sensitivity functions corresponding to these three parameters also exhibit linear dependence (Figure 5.C).

Among the free parameters estimated in the calibration completion stage (Figure 5.D), the largest sensitivities are observed for Glc, particularly for parameters associated with adsorption of cellulase enzymes on cellulose, namely 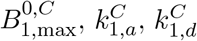. The sensitivity functions of Glc with respect to 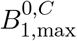 and 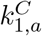 are nearly proportional to the sensitivity function associated with the stage-2 parameter *k*_*r,C*_, indicating a similar influence of these parameters on the predicted Glc conversion dynamics. The parameters 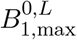 and 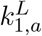 have a moderate influence on Glc conversion dynamics. Similarly, the parameters 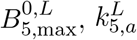, and 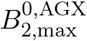 have a moderate influence on the Xyl conversion dynamics. Nearly all of the remaining calibration completion stage parameters (Figure 5.D) have some influence on the dynamics of Glc and Xyl.

A first implication of the sensitivity analysis is that, in each of identification stages 1, 2, and 3, the free parameters exert a non-negligible influence primarily on the target variable associated with the dominant branch defined for that stage (see Table S1 in the supplementary material). Specifically, the free parameters of stages 1, 2, and 3 predominantly affect Man, Glc, and Xyl, respectively. This behavior is consistent with the stage-and-sugar specific weights in the objective function. The sensitivity results therefore support the rationale underlying the staged estimation procedure and validate the structure of the proposed estimation method.

The sensitivity analysis also showed that nearly all parameters simplified in the transition from DyAIM to ReAIM and EAM exert a negligible influence on the target variables. In particular, among the 19 simplified parameters, 14 provide normalized sensitivities with a maximal absolute value below 2%, for all the target variables (and even below 1% for 12 of these parameters). The 5 remaining simplified parameters are among the parameters simplified in the transition from ReAIM to EAM, namely 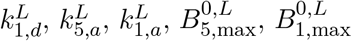. They exhibit higher, still moderate, sensitivities, with maximum absolute value ranging from 4.4% to 10.1% (resp. from 2.6% to 5.7%) for the enzyme loading of 15 FPU/g biomass (resp. of 30 FPU/g biomass). The sensitivity analysis thus provides complementary support for and further reinforces the mechanistic model simplifications adopted in ReAIM and EAM.

### 3.8 The hierarchy offers an integrative mechanistic atlas of lignocellulosic hydrolysis

Beyond its predictive role, the model hierarchy developed in this study can be viewed as an integrative mechanistic atlas of enzyme–cell-wall interactions. It organizes adsorption relationships between enzyme pools and structural polymers, including cellulose, hemicelluloses, and lignin, together with the inhibitory effects of hydrolysis products, into a coherent framework. By progressively increasing mechanistic details from EAM to ReAIM and DyAIM, the hierarchy represents these coupled processes at different levels of detail and provides a structured representation of the mechanisms shaping hydrolysis dynamics. Thus, rather than constituting only a family of kinetic models, the hierarchy also serves as an integrative synthesis of current mechanistic knowledge on enzyme adsorption and product inhibition during lignocellulosic hydrolysis, by integrating dispersed mechanistic evidence into a unified model architecture.

This framework also provides a foundation for incorporating additional scales representing enzymatic hydrolysis. One complementary extension would be to explicitly represent enzyme transport and molecular-scale interactions. Enzyme diffusion and penetration within the cell wall could, for instance, be parameterized using diffusivities and mobility estimates derived from fluorescence recovery after photobleaching (FRAP) measurements, linking local substrate accessibility and enzyme mobility to effective catalytic activity [19, 48]. Such developments would provide a more mechanistic description of how enzyme transport, confinement, and interfacial dynamics contribute to the observed hydrolysis kinetics.

A further extension would be to incorporate cell- and tissue-scale hydrolysis dynamics extracted from four-dimensional (3D space + time) imaging of plant cell walls [20, 47, 70]. Integrating such spatially and temporally resolved information would enable the framework to account explicitly for heterogeneity in cell-wall architecture, and the progression of deconstruction across cells and tissues. Together, these extensions could move the present hierarchy toward a genuinely multiscale description of lignocellulosic hydrolysis, linking molecular-scale enzyme interactions and transport processes to cell-wall structure, tissue organization, and macroscopic conversion dynamics.

## 4 Conclusions

This work establishes a hierarchical adsorption–inhibition modeling framework for lignocellulosic enzymatic hydrolysis that links mechanistic enzyme–substrate interactions to sugar-release dynamics. The framework comprises three nested formulations with progressively reduced complexity: the comprehensive Dynamic Adsorption–Inhibition Model (DyAIM), the Reduced Adsorption–Inhibition Model (ReAIM), in which weak adsorption and inhibition interactions are removed, and the Effective Activity Model (EAM), in which nonproductive lignin adsorption is represented through a static reduction in effective enzyme activity. This hierarchy therefore combines mechanistic resolution with systematic model reduction, resulting in increasingly compact yet mechanistically consistent descriptions of enzymatic hydrolysis.

All three models accurately reproduced glucose, xylose, and mannose release across different enzyme loadings and retained robust predictive performance during external validation at an unseen loading of 7.5 FPU/g biomass. The fitted inhibition constants captured relative product-inhibition strengths consistent with literature trends, further supporting the mechanistic basis of the framework. Sensitivity analysis provided complementary support for the structural reductions adopted in ReAIM and EAM. Beyond prediction, the hierarchy also forms a coherent mechanistic atlas of the processes governing lignocellulosic hydrolysis.

Overall, the comparable predictive performance of progressively simplified formulations demonstrates that nonessential complexity can be removed without compromising the dominant system behavior. The proposed hierarchy therefore provides a systematic basis for distinguishing essential mechanisms from dispensable complexity and for selecting an appropriate level of model detail. Beyond mechanistic interpretation, the framework offers a basis for process optimization, enzyme-cocktail design, and decision-making in lignocellulosic biomass conversion, and could support future development of data-integrated digital-twin applications.

## CRediT authorship contribution statement

**Solmaz Hossein Khani:** Conceptualization, Data curation, Formal analysis, Investigation, Methodology, Software, Validation, Visualization, Writing – original draft, Writing – review and editing. **Ali Faraj:** Methodology, Software, Writing – original draft, Writing – review and editing. **Grégoire Malandain:** Investigation, Methodology, Writing – review and editing. **Gabriel Paës:** Conceptualization, Formal analysis, Funding acquisition, Investigation, Methodology, Project administration, Resources, Supervision, Validation, Writing – review and editing. **Yassin Refahi:** Conceptualization, Data curation, Formal analysis, Funding acquisition, Investigation, Methodology, Project administration, Resources, Software, Supervision, Validation, Visualization, Writing – original draft, Writing – review and editing.

## Supplementary material

This article is accompanied by supplementary material containing additional information.

## Acknowledgments

The authors gratefully acknowledge Simon Arragain and Antoine Margeot (IFPEN, Rueil-Malmaison, France) for providing the enzymatic cocktail. They also acknowledge Juliette Floret for conducting the enzymatic activity assays, François Gaudard, Anouck Habrant, Noah Remy, and Berangère Lebas for sample preparation and conversion-yield measurements.

## Funding acknowledgments

This work was funded by French Research Minister aid managed by the Agence Nationale de la Recherche as part of the France 2030 investment plan through the grant reference ANR-23-PEBB-0006 (FillingGaps project). This work was also funded by Agence Nationale de la Recherche (ANR) under grant ANR-19-CE43-0010 (BIOMOD project), and by Grand Est Region through “BIOMODEL” doctoral funding.

