## Supplementary Material for "Taming complexity in enzymatic saccharification: A predictive hierarchical modeling framework"

#### Contents

**S1 Calculation of polymer-class mass fractions** **2**

**S2 Enzyme activity assignments** **2**

**S3 Explicit equations of the adsorption kinetics in DyAIM** **2**

**S4 Explicit adsorption and enzyme activity concentration balance equations in ReAIM** **3**

**S5 Explicit productive adsorption equations in EAM** **4**

**S6 Multi-stage parameter estimation procedure** **5**

**S7 Fit-quality assessment** **9**

**S8 Implementation of the models and the parameter estimation** **12**

**S9 Parametric sensitivity analysis** **13**

### S1 Calculation of polymer-class mass fractions

The polymer-class mass fractions, expressed as mass percentages of the original pretreated sample, were calculated as follows with $w$ denoting the corresponding sugar mass fraction: cellulose fraction prior to hydrolysis, $w^0_{\mathrm{C}}$, was estimated as $w_{\mathrm{Glc}} - \dfrac{w_{\mathrm{Man}}}{4}$, whereas GGM fraction prior to hydrolysis, $w^0_{\mathrm{GGM}}$, was calculated as $w_{\mathrm{Man}} + w_{\mathrm{Gal}} + \dfrac{w_{\mathrm{Man}}}{4}$. Initial AGX fraction, $w^0_{\mathrm{AGX}}$, was defined as $w_{\mathrm{Xyl}} + w_{\mathrm{Ara}} + w_{\mathrm{GlcA}}$, and the initial lignin fraction, $w^0_{\mathrm{L}}$, was taken directly from the measured lignin content.

### S2 Enzyme activity assignments

The measured cellulase, xylanase, endo-1,4-$\beta$-mannanase, $\beta$-glucosidase, and $\beta$-xylosidase activities were assigned to pools $\mathrm{E}_1$, $\mathrm{E}_2$, $\mathrm{E}_3$, $\mathrm{E}_4$, and $\mathrm{E}_5$, respectively. These assignments indicate lumped functional pools. The $\alpha$-arabinosidase and acetyl esterase were not represented as independent pools; instead, their contributions were implicitly incorporated into $\mathrm{E}_2$. Similarly, the measured endo-1,4-$\beta$-mannanase activity was used as the experimental proxy for the composite mannanolytic pool $\mathrm{E}_3$, which represents both GGM depolymerization and subsequent conversion of MOS in the lumped model.

### S3 Explicit equations of the adsorption kinetics in DyAIM

The adsorption dynamics in the kinetic model presented in the main manuscript are based on a dynamic Langmuir kinetics, in which each enzyme pool may bind either productively or nonproductively to a given component. Specifically, in DyAIM, $\mathrm{E}_1$ is assumed to bind productively to cellulose and nonproductively to lignin. $\mathrm{E}_2$ is assumed to bind productively to AGX and nonproductively to lignin. The mannanolytic pool $\mathrm{E}_3$ is assumed to bind productively to GGM (with nonproductive binding to lignin neglected). The soluble glycosidase pools $\mathrm{E}_4$ and $\mathrm{E}_5$ are not assumed to bind productively to insoluble polysaccharides, but they may be depleted from solution by nonproductive adsorption to lignin. The main text introduced the general formulation and the fractional occupancy terms denoted by $\chi^s(t)$, for component $s$.

For cellulose, AGX, and GGM, the fractional occupancies are defined, respectively, as

$$\chi^C(t) = \frac{b_1^C(t)}{B_{1,\max}^C(t)}, \qquad \chi^{\mathrm{AGX}}(t) = \frac{b_2^{\mathrm{AGX}}(t)}{B_{2,\max}^{\mathrm{AGX}}(t)}, \qquad \chi^{\mathrm{GGM}}(t) = \frac{b_3^{\mathrm{GGM}}(t)}{B_{3,\max}^{\mathrm{GGM}}(t)}.$$

Using these definitions of the fractional occupancies, the explicit equation corresponding to $b_1^C(t)$, $b_1^L(t)$, $b_2^{\mathrm{AGX}}(t)$, $b_2^L(t)$, $b_3^{\mathrm{GGM}}(t)$, $b_4^L(t)$, and $b_5^L(t)$ are provided below:

$$\frac{db_1^C}{dt} = k_{1,a}^C\, E_{1,f}(t)\, B_{1,\max}^C(t) \left(1 - \frac{b_1^C(t)}{B_{1,\max}^C(t)}\right) - k_{1,d}^C\, b_1^C(t),$$

$$\frac{db_1^L}{dt} = k_{1,a}^L\, E_{1,f}(t)\, B_{1,\max}^L(t) \left(1 - \chi^L(t)\right) - k_{1,d}^L\, b_1^L(t),$$

$$\frac{db_2^{\mathrm{AGX}}}{dt} = k_{2,a}^{\mathrm{AGX}}\, E_{2,f}(t)\, B_{2,\max}^{\mathrm{AGX}}(t) \left(1 - \frac{b_2^{\mathrm{AGX}}(t)}{B_{2,\max}^{\mathrm{AGX}}(t)}\right) - k_{2,d}^{\mathrm{AGX}}\, b_2^{\mathrm{AGX}}(t),$$

$$\frac{db_2^L}{dt} = k_{2,a}^L\, E_{2,f}(t)\, B_{2,\max}^L(t)\left(1 - \chi^L(t)\right) - k_{2,d}^L\, b_2^L(t),$$

$$\frac{db_3^{\mathrm{GGM}}}{dt} = k_{3,a}^{\mathrm{GGM}}\, E_{3,f}(t)\, B_{3,\max}^{\mathrm{GGM}}(t)\left(1 - \frac{b_3^{\mathrm{GGM}}(t)}{B_{3,\max}^{\mathrm{GGM}}(t)}\right) - k_{3,d}^{\mathrm{GGM}}\, b_3^{\mathrm{GGM}}(t),$$

$$\frac{db_4^L}{dt} = k_{4,a}^L\, E_{4,f}(t)\, B_{4,\max}^L(t)\left(1 - \chi^L(t)\right) - k_{4,d}^L\, b_4^L(t),$$

$$\frac{db_5^L}{dt} = k_{5,a}^L\, E_{5,f}(t)\, B_{5,\max}^L(t)\left(1 - \chi^L(t)\right) - k_{5,d}^L\, b_5^L(t),$$

where $\chi^L(t)$ is the shared lignin occupancy as defined in the main text:

$$\chi^L(t) = \frac{b_1^L(t)}{B_{1,\max}^L(t)} + \frac{b_2^L(t)}{B_{2,\max}^L(t)} + \frac{b_4^L(t)}{B_{4,\max}^L(t)} + \frac{b_5^L(t)}{B_{5,\max}^L(t)}.$$

Furthermore, with

$$\rho_C(t) = C(t), \qquad \rho_{\mathrm{AGX}}(t) = H_{\mathrm{AGX}}(t), \qquad \rho_{\mathrm{GGM}}(t) = H_{\mathrm{GGM}}(t), \qquad \rho_L(t) = L(t),$$

the enzyme activity concentration balances are given by

$$\begin{aligned}
E_{1,\mathrm{tot}} &= E_{1,f}(t) + b_1^C(t)\, C(t) + b_1^L(t)\, L(t),\\
E_{2,\mathrm{tot}} &= E_{2,f}(t) + \; b_2^{\mathrm{AGX}}(t)\, H_{\mathrm{AGX}}(t) + b_2^L(t) L(t),\\
E_{3,\mathrm{tot}} &= E_{3,f}(t) + b_3^{\mathrm{GGM}}(t)\, H_{\mathrm{GGM}}(t),\\
E_{4,\mathrm{tot}} &= E_{4,f}(t) + b_4^L(t)\, L(t),\\
E_{5,\mathrm{tot}} &= E_{5,f}(t) + b_5^L(t)\, L(t).
\end{aligned}$$

At $t = 0$, no enzyme is assumed to be bound, so that $b_i^s(0) = 0$ for all admissible $(i, s)$ pairs, and therefore $E_{i,f}(0) = E_{i,\mathrm{tot}}$.

### S4 Explicit adsorption and enzyme activity concentration balance equations in ReAIM

In ReAIM, the binding components sets are

$$\mathcal{S}_1^{\mathrm{bind}} = \{C, L\}, \quad \mathcal{S}_2^{\mathrm{bind}} = \{\mathrm{AGX}\}, \quad \mathcal{S}_3^{\mathrm{bind}} = \{\mathrm{GGM}\}, \quad \mathcal{S}_4^{\mathrm{bind}} = \{L\}, \quad \mathcal{S}_5^{\mathrm{bind}} = \{L\}.$$

Compared with DyAIM, ReAIM omits the nonproductive binding of $E_2$ to lignin and fixes all maximum adsorption capacities to the constant baseline adsorption capacities

$$B_{1,\max}^{0,C},\; B_{2,\max}^{0,\mathrm{AGX}},\; B_{3,\max}^{0,\mathrm{GGM}},\; B_{i,\max}^{0,L}\; (i = 1, 4, 5).$$

Dynamic adsorption is described using dynamic Langmuir kinetics as in DyAIM. By defining the shared lignin occupancy as

$$\chi^L(t) = \frac{b_1^L(t)}{B_{1,\max}^{0,L}} + \frac{b_4^L(t)}{B_{4,\max}^{0,L}} + \frac{b_5^L(t)}{B_{5,\max}^{0,L}}, \qquad 0 \le \chi^L(t) \le 1,$$

the equations describing adsorption are:

$$\begin{aligned}
\frac{db_1^C}{dt} &= k_{1,a}^C\, E_{1,f}(t)\, B_{1,\max}^{0,C}\Big(1 - \frac{b_1^C(t)}{B_{1,\max}^{0,C}}\Big) - k_{1,d}^C\, b_1^C(t),\\
\frac{db_1^L}{dt} &= k_{1,a}^L\, E_{1,f}(t)\, B_{1,\max}^{0,L}\big(1 - \chi^L(t)\big) - k_{1,d}^L\, b_1^L(t),\\
\frac{db_2^{\mathrm{AGX}}}{dt} &= k_{2,a}^{\mathrm{AGX}}\, E_{2,f}(t)\, B_{2,\max}^{0,\mathrm{AGX}}\Big(1 - \frac{b_2^{\mathrm{AGX}}(t)}{B_{2,\max}^{0,\mathrm{AGX}}}\Big) - k_{2,d}^{\mathrm{AGX}}\, b_2^{\mathrm{AGX}}(t),\\
\frac{db_3^{\mathrm{GGM}}}{dt} &= k_{3,a}^{\mathrm{GGM}}\, E_{3,f}(t)\, B_{3,\max}^{0,\mathrm{GGM}}\Big(1 - \frac{b_3^{\mathrm{GGM}}(t)}{B_{3,\max}^{0,\mathrm{GGM}}}\Big) - k_{3,d}^{\mathrm{GGM}}\, b_3^{\mathrm{GGM}}(t),\\
\frac{db_4^L}{dt} &= k_{4,a}^L\, E_{4,f}(t)\, B_{4,\max}^{0,L}\big(1 - \chi^L(t)\big) - k_{4,d}^L\, b_4^L(t),\\
\frac{db_5^L}{dt} &= k_{5,a}^L\, E_{5,f}(t)\, B_{5,\max}^{0,L}\big(1 - \chi^L(t)\big) - k_{5,d}^L\, b_5^L(t).
\end{aligned}$$

The enzyme activity concentration balances are

$$\begin{aligned}
E_{1,\mathrm{tot}} &= E_{1,f}(t) + b_1^C(t)\, C(t) + b_1^L(t)\, L(t),\\
E_{2,\mathrm{tot}} &= E_{2,f}(t) + b_2^{\mathrm{AGX}}(t)\, H_{\mathrm{AGX}}(t),\\
E_{3,\mathrm{tot}} &= E_{3,f}(t) + b_3^{\mathrm{GGM}}(t)\, H_{\mathrm{GGM}}(t),\\
E_{4,\mathrm{tot}} &= E_{4,f}(t) + b_4^L(t)\, L(t),\\
E_{5,\mathrm{tot}} &= E_{5,f}(t) + b_5^L(t)\, L(t).
\end{aligned}$$

Similar to DyAIM, at $t = 0$, no enzyme is assumed to be bound, so that $b_i^s(0) = 0$ for all admissible enzyme–surface pairs $(i, s)$, and therefore $E_{i,f}(0) = E_{i,\mathrm{tot}}$.

### S5 Explicit productive adsorption equations in EAM

In EAM, the effective catalytic activity concentrations are defined as

$$E_{i,\,\mathrm{eff}} = (1 - \phi_i)\, E_{i,\,\mathrm{tot}},\; i \in \{1, 4, 5\},$$

with $0 \leq \phi_i \leq 1$. No lignin-adsorption correction is applied to $E_2$ and $E_3$ (i.e., $\phi_2 = \phi_3 = 0$), thus $E_{i,\,\text{eff}} = E_{i,\,\text{tot}}$, $i \in \{2, 3\}$. $E_{i,\text{eff}}$ replaces $E_{i,\text{tot}}$ to compute the free enzyme activity concentrations:

$$E_{1,f}(t) = E_{1,\text{eff}} - b_1^C(t)\,C(t)\,, \qquad E_{2,f}(t) = E_{2,\text{eff}} - b_2^{\text{AGX}}(t)\,H_{\text{AGX}}(t)\,,$$

$$E_{3,f}(t) = E_{3,\text{eff}} - b_3^{\text{GGM}}(t)\,H_{\text{GGM}}(t)\,, \quad E_{4,f}(t) = E_{4,\text{eff}}\,, \qquad E_{5,f}(t) = E_{5,\text{eff}}\,.$$

Dynamic adsorption is restricted to productive binding on polysaccharides. Therefore, compared to DyAIM and ReAIM, only $b_1^C$, $b_2^{\text{AGX}}$, and $b_3^{\text{GGM}}$ are retained with the following adsorption equations:

$$\frac{db_1^C}{dt} = k_{1,a}^C\,E_{1,f}(t)\,B_{1,\max}^{0,C}\Big(1 - \frac{b_1^C(t)}{B_{1,\max}^{0,C}}\Big) - k_{1,d}^C\,b_1^C(t),$$

$$\frac{db_2^{\text{AGX}}}{dt} = k_{2,a}^{\text{AGX}}\,E_{2,f}(t)\,B_{2,\max}^{0,\,\text{AGX}}\Big(1 - \frac{b_2^{\text{AGX}}(t)}{B_{2,\max}^{0,\text{AGX}}}\Big) - k_{2,d}^{\text{AGX}}\,b_2^{\text{AGX}}(t),$$

$$\frac{db_3^{\text{GGM}}}{dt} = k_{3,a}^{\text{GGM}}\,E_{3,f}(t)\,B_{3,\max}^{0,\,\text{GGM}}\Big(1 - \frac{b_3^{\text{GGM}}(t)}{B_{3,\max}^{0,\,\text{GGM}}}\Big) - k_{3,d}^{\text{GGM}}\,b_3^{\text{GGM}}(t).$$

At $t = 0$, no enzymes are bound to polysaccharides, thus $b_1^C(0) = b_2^{\text{AGX}}(0) = b_3^{\text{GGM}}(0) = 0$, and $E_{i,f}(0) = E_{i,\text{eff}}$, $1 \leq i \leq 5$.

### S6 Multi-stage parameter estimation procedure

The multi-stage parameter estimation procedure for a given model involved four consecutive parameter-estimation stages. At each stage, only a predefined subset of parameters influencing the stage-specific dominant data was allowed to vary. The parameter estimates obtained at stage $s \in \{1, 2, 3\}$ were used in stage $s + 1$, propagating information through the successive stages. Parameter estimation at each stage $s$ was formulated as a nonlinear least-squares problem. For a target sugar $j \in$ {Glc, Xyl, Man}, enzyme loading $f \in \{15, 30\}$, and sampling time $t_i$, the weighted residual was defined as

$$r_{j,f}^s(t_i; \boldsymbol{\theta}) = \omega_j^s\,[\hat{y}_{j,f}(t_i; \boldsymbol{\theta}) - y_{j,f}(t_i)]\,,$$

where $y_{j,f}(t_i)$ denotes the experimentally measured conversion of sugar $j$ at enzyme loading $f$ and sampling time $t_i$, and $\hat{y}_{j,f}(t_i; \boldsymbol{\theta})$ denotes the predicted conversion yield at sampling time $t_i$ for the parameter vector $\boldsymbol{\theta}$. The coefficient $\omega_j^s \geq 0$ is a stage- and sugar-specific weight.

- Stage 1. Only parameters influencing Man yield were optimized, based on Man conversion yields at both enzyme loadings. Accordingly, the Man residual was assigned a weight $\omega_{\text{Man}}^1 = 1.0$, while Glc and Xyl residuals were set to zero.

- Stage 2. Only parameters affecting Glc yield were estimated, with Glc conversion yields at both enzyme loadings as the dominant observables. The Glc residuals were assigned the largest weight ($\omega_{\text{Glc}}^2 = 1.0$), Man residuals a moderate weight ($\omega_{\text{Man}}^2 = 0.5$), and Xyl residuals were set to zero. This weighting drives the estimation of Glc-related parameters while preventing large deviations in

Table S1: Free parameters and sugar-specific weights in multi-stage parameter estimation procedure. For each model (DyAIM, ReAIM, EAM) only the parameters present in that model are treated as free in a given stage; parameters eliminated by simplification are omitted automatically.

| Stage | Dominant sugar | Free parameters | Weights |
|---|---|---|---|
| **1** | Man – 15 & 30 FPU | $B_{3,\max}^{0,\mathrm{GGM}}, k_{3,a}^{\mathrm{GGM}}, k_{3,d}^{\mathrm{GGM}}, k_{r,\mathrm{GGM}}, k_{\mathrm{cat},3}, K_{M,3}, K_{I,\mathrm{GGM}}^{M_1}, K_{I,3}^{M_1}$ | $\omega_{\mathrm{Man}}^1 = 1.0$, $\omega_{\mathrm{Glc}}^1 = 0.0$, $\omega_{\mathrm{Xyl}}^1 = 0.0$ |
| **2** | Glc – 15 & 30 FPU | $k_{r,C}, k_{\mathrm{cat},4}, K_{M,4}, K_{I,4}^{G_1}, K_{I,4}^{G_2}, K_{I,C}^{G_2}, K_{I,C}^{G_1}, K_{I,C}^{\mathrm{XOS}}, K_{I,C}^{X_1}, K_{I,C}^{M_1}, K_{I,C}^{\mathrm{MOS}}, \alpha$ | $\omega_{\mathrm{Man}}^2 = 0.5$, $\omega_{\mathrm{Glc}}^2 = 1.0$, $\omega_{\mathrm{Xyl}}^2 = 0.0$ |
| **3** | Xyl – 15 & 30 FPU | $k_{r,\mathrm{AGX}}, k_{\mathrm{cat},5}, K_{M,5}, K_{I,5}^{X_1}, K_{I,\mathrm{AGX}}^{\mathrm{XOS}}, K_{I,\mathrm{AGX}}^{G_2}, K_{I,\mathrm{AGX}}^{G_1}, K_{I,\mathrm{AGX}}^{\mathrm{MOS}}, k_{2,a}^{\mathrm{AGX}}, k_{2,d}^{\mathrm{AGX}}$ | $\omega_{\mathrm{Man}}^3 = 0.5$, $\omega_{\mathrm{Glc}}^3 = 0.1$, $\omega_{\mathrm{Xyl}}^3 = 1.0$ |
| **4** | All three sugars – 15 & 30 FPU | *Parameters not estimated in Stages 1–3* | $\omega_{\mathrm{Man}}^4 = 1.0$, $\omega_{\mathrm{Glc}}^4 = 1.0$, $\omega_{\mathrm{Xyl}}^4 = 1.0$ |

the previously fitted Man trajectories.

- Stage 3. Only parameters associated with Xyl yield were optimized, with Xyl conversion yields at both enzyme loadings as the primary observables. The Xyl residual received full weight $\omega_{\mathrm{Xyl}}^3 = 1.0$, while Glc and Man residuals were given lower weights ($\omega_{\mathrm{Glc}}^3 = 0.1$, $\omega_{\mathrm{Man}}^3 = 0.5$) to act as soft constraints that preserve the fits obtained in the earlier stages.
- Stage 4. All parameters not estimated in Stages 1–3 were fitted, assigning equal weights $\omega_{\mathrm{Glc}}^4 = \omega_{\mathrm{Man}}^4 = \omega_{\mathrm{Xyl}}^4 = 1.0$.

For a given model, the objective function minimized at stage $s$ was therefore

$$\Phi^s(\boldsymbol{\theta}) = \sum_{j\in\{\mathrm{Glc,\,Xyl,\,Man}\}} \sum_{f\in\{15,30\}} \sum_{t_i\in\mathcal{T}} \left[r_{j,f}^s(t_i;\boldsymbol{\theta})\right]^2,$$

where $\mathcal{T}$ denotes the set of sampling times.

Details of the numerical implementation of the models and the parameter estimation procedure are provided in Section S8.

Initial parameter values for the EAM, as well as the parameters not shared across models (i.e., not transferred from simpler models to more detailed models in warm-start strategy) were set based on values reported for similar systems in the literature and on expert judgment. For each free parameter, lower and upper bounds were also specified from values reported for similar systems in the literature and on expert judgment. See Table S1 for free parameters and sugar-specific weights at each stage.

In EAM, enzyme-specific lignin adsorption factors were taken from adsorption measurements of the major *Trichoderma reesei* enzymes on pine lignin, with pine used as a softwood analog for spruce [3]. For $E_i$, $i \in \{1,4,5\}$, the corresponding fraction adsorbed on lignin is scaled by the lignin content in the pretreated sample (11.97%) which yielded $\phi_1 = 0.0088652$, $\phi_4 = 0.0679$, $\phi_5 = 0.00243$.

Table S2: Experimentally determined parameters. These values are common to DyAIM, ReAIM, and EAM.

| Symbol | Definition | Value |
|---|---|---|
| $C^0$ | Initial cellulose concentration (mg/mL) | 4.217129 |
| $H^0_{AGX}$ | Initial arabino-4-$O$-methyl-glucuronoxylan (AGX) concentration (mg/mL) | 0.584731 |
| $H^0_{GGM}$ | Initial galactoglucomannan (GGM) concentration (mg/mL) | 1.544382 |
| $L^0$ | Initial lignin concentration (mg/mL) | 0.863758 |
| $G^0_1$ | Initial Glc concentration (mg/mL) | 0 |
| $G^0_2$ | Initial $G_2$ concentration (mg/mL) | 0 |
| $X^0_1$ | Initial Xyl concentration (mg/mL) | 0 |
| $M^0_1$ | Initial Man concentration (mg/mL) | 0 |
| $XOS^0$ | Initial XOS concentration (mg/mL) | 0 |
| $MOS^0$ | Initial MOS concentration (mg/mL) | 0 |
| $A_1$ | Measured cellulase activity (filter paper assay) (FPU/mL) | 2.7 |
| $A_2$ | Measured xylanase activity (Kidby assay) (IU/mL) | 9.33 |
| $A_3$ | Measured endo-1,4-$\beta$-mannanase activity on konjac glucomannan (IU/mL) | 0.94 |
| $A_4$ | Measured $\beta$-glucosidase activity (pNP-glucopyranoside assay) (IU/mL) | 5.1 |
| $A_5$ | Measured $\beta$-xylosidase activity (pNP-xylopyranoside assay) (IU/mL) | 3.25 |
| $\eta$ | Enzyme cocktail dose factor (mL enzyme cocktail per mL slurry) for the 7.5, 15, and 30 FPU/g biomass | 0.0245/0.049/0.098 |

Table S3: Optimal parameter values for DyAIM, ReAIM, and EAM.

| Parameter | DyAIM | ReAIM | EAM | Parameter | DyAIM | ReAIM | EAM |
|---|---|---|---|---|---|---|---|
| $\alpha$ | 2.480 | 2.519 | 2.395 | $\gamma_C$ | 0.02571 | – | – |
| $\gamma_L$ | 0.6529 | – | – | $B^{0,C}_{1,\max}$ | 0.1523 | 0.1308 | 0.07449 |
| $B^{0,\mathrm{AGX}}_{2,\max}$ | 0.8000 | 0.7656 | 0.7408 | $B^{0,\mathrm{GGM}}_{3,\max}$ | 0.03000 | 0.03000 | 0.03000 |
| $B^{0,L}_{1,\max}$ | 0.3674 | 0.3543 | – | $B^{0,L}_{2,\max}$ | 0.7669 | – | – |
| $B^{0,L}_{4,\max}$ | 0.2797 | 0.04388 | – | $B^{0,L}_{5,\max}$ | 0.2900 | 0.1877 | – |
| $k^C_{1,a}$ | 0.5312 | 0.5811 | 0.7546 | $k^C_{1,d}$ | 0.6906 | 0.6632 | 1.0000 |
| $k^L_{1,a}$ | 1.454 | 1.395 | – | $k^L_{1,d}$ | 0.07632 | 0.05989 | – |
| $k^{\mathrm{AGX}}_{2,a}$ | 0.4927 | 0.4953 | 0.4974 | $k^{\mathrm{AGX}}_{2,d}$ | 0.5488 | 0.06264 | 0.06261 |
| $k^L_{2,a}$ | 0.2993 | – | – | $k^L_{2,d}$ | 0.3668 | – | – |
| $k^{\mathrm{GGM}}_{3,a}$ | 0.2051 | 1.338 | 1.543 | $k^{\mathrm{GGM}}_{3,d}$ | 1.0000 | 1.0000 | 1.0000 |
| $k^L_{4,a}$ | 0.01119 | 0.0005531 | – | $k^L_{4,d}$ | 0.2610 | 0.04330 | – |
| $k^L_{5,a}$ | 0.1393 | 0.1245 | – | $k^L_{5,d}$ | 0.005002 | 0.005040 | – |
| $k_{r,C}$ | 199.9 | 199.9 | 199.9 | $k_{r,\mathrm{AGX}}$ | 290.2 | 289.8 | 289.8 |
| $k_{r,\mathrm{GGM}}$ | 36.90 | 5.684 | 4.952 | $K^{\mathrm{G}_2}_{I,C}$ | 5.955 | 6.869 | 11.83 |
| $K^{\mathrm{G}_1}_{I,C}$ | 62.65 | – | – | $K^{\mathrm{XOS}}_{I,C}$ | 4.480 | 5.994 | 5.970 |
| $K^{\mathrm{X}_1}_{I,C}$ | 89.07 | – | – | $K^{\mathrm{M}_1}_{I,C}$ | 93.22 | – | – |
| $K^{\mathrm{MOS}}_{I,C}$ | 11.43 | 11.53 | 11.57 | $K^{\mathrm{XOS}}_{I,\mathrm{AGX}}$ | 2.136 | 2.520 | 2.518 |
| $K^{\mathrm{G}_2}_{I,\mathrm{AGX}}$ | 51.26 | 21.97 | 21.96 | $K^{\mathrm{G}_1}_{I,\mathrm{AGX}}$ | 151.9 | – | – |
| $K^{\mathrm{MOS}}_{I,\mathrm{AGX}}$ | 10.01 | 4.458 | 4.458 | $K^{\mathrm{M}_1}_{I,\mathrm{GGM}}$ | 0.6657 | 1.418 | 1.654 |
| $k_{\mathrm{cat},4}$ | 24.98 | 24.73 | 24.82 | $K_{M,4}$ | 0.1000 | 0.1000 | 0.1000 |
| $K^{\mathrm{G}_1}_{I,4}$ | 91.29 | 94.56 | 62.05 | $K^{\mathrm{G}_2}_{I,4}$ | 96.33 | – | – |
| $k_{\mathrm{cat},5}$ | 13.64 | 12.35 | 12.35 | $K_{M,5}$ | 8.589 | 8.305 | 8.305 |
| $K^{\mathrm{X}_1}_{I,5}$ | 0.1000 | 0.1000 | 0.1000 | $k_{\mathrm{cat},3}$ | 25.00 | 25.00 | 25.00 |
| $K_{M,3}$ | 0.03000 | 0.03000 | 0.03000 | $K^{\mathrm{M}_1}_{I,3}$ | 2.788 | 0.8763 | 7.315 |

#### S7 Fit-quality assessment

Model fit quality was assessed using the range-normalized root mean square error (NRMSE) and the coefficient of determination, $R^2$. The NRMSE was defined as the RMSE divided by the observed range of the corresponding sugar–enzyme-loading block and expressed as a percentage of that range. Metrics were computed for each sugar at each enzyme loading, globally by pooling the three sugars at each enzyme loading, and also globally by pooling the three sugars across both enzyme loadings. In other words, for a given model, pooling involved either both enzyme loadings (15 and 30 FPU/g biomass) or all sugars at a given enzyme loading. To calculate global NRMSE values, residuals were normalized by the range of their respective sugar–enzyme-loading block to account for differences in sugar conversion yield magnitude. For pooled datasets, the reported global $R^2$ was calculated as the mean of the $R^2$ values for the constituent individual-sugar datasets.

More precisely, for a given model, these metrics were computed for each enzyme loading $f \in \{15, 30\}$ FPU/g biomass, and each target sugar $j \in \{\text{Glc, Xyl, Man}\}$. As defined in Section S6, let $\mathcal{T}$ denote the common set of sampling times, and let $y_{j,f}(t_i)$ denote the experimentally measured conversion of sugar $j$ at enzyme loading $f$ and sampling time $t_i$. The corresponding prediction evaluated at the parameter vector $\boldsymbol{\theta}$, is denoted by $\hat{y}_{j,f}(t_i;\boldsymbol{\theta})$. The unweighted residual used to compute the fit-quality metrics was defined as

$$\varepsilon_{j,f}(t_i) = \hat{y}_{j,f}(t_i;\boldsymbol{\theta}) - y_{j,f}(t_i).$$

For enzyme loading $f$ , and target sugar $j$, the root mean square error was computed as

$$\text{RMSE}_{f,j} = \left[\frac{1}{|\mathcal{T}|}\sum_{t_i\in\mathcal{T}} \left(\varepsilon_{j,f}(t_i)\right)^2\right]^{1/2},$$

where $|\mathcal{T}|$ is the number of sampling times, which was identical for all target sugars and both enzyme loadings.

Because sugar conversion yields differ in magnitude and range, raw RMSE values do not provide a fair basis for comparing relative fit quality across sugars. A given absolute error represents a much larger proportion of the mannose conversion range than it does of the Glc or Xyl conversion yield ranges. Range normalization is therefore useful when comparing model performance among sugars and is essential before pooling them into a single global metric, otherwise the larger-range Glc and Xyl conversions would contribute disproportionately. To enable comparisons among target sugars with different observed dynamic ranges, the RMSE was normalized by the observed range of the corresponding sugar–enzyme-loading data block. For sugar $j$ at enzyme loading $f$, this range was defined as

$$\Delta y_{j,f} = \max_{t_i\in\mathcal{T}} y_{j,f}(t_i) - \min_{t_i\in\mathcal{T}} y_{j,f}(t_i).$$

The sugar-specific range-normalized RMSE expressed as a percentage the observed range of conversion yield of sugar $j$ at enzyme loading $f$ was then computed as

$$\text{NRMSE}_{f,j} = 100\frac{\text{RMSE}_{f,j}}{\Delta y_{j,f}}.$$

The sugar-specific coefficient of determination was computed as

$$R^2_{f,j} = 1 - \frac{\sum_{t_i \in \mathcal{T}} \left(\varepsilon_{j,f}(t_i)\right)^2}{\sum_{t_i \in \mathcal{T}} \left(y_{j,f}(t_i) - \bar{y}_{j,f}\right)^2},$$

where

$$\bar{y}_{j,f} = \frac{1}{|\mathcal{T}|} \sum_{t_i \in \mathcal{T}} y_{j,f}(t_i)$$

is the mean observed conversion of sugar $j$ at enzyme loading $f$.

In addition to the sugar-specific metrics, global fit-quality metrics were computed by pooling the residuals across the three target sugars. As mentioned previously, the reported global $R^2$ was calculated as the mean of the $R^2$ values for the three target sugars datasets. For the global NRMSE, the residuals of each sugar were first normalized separately within each enzyme-loading level by the observed range of the corresponding sugar–enzyme-loading data block. The normalized residual was defined as

$$\widetilde{\varepsilon}_{j,f}(t_i) = \frac{\varepsilon_{j,f}(t_i)}{\Delta y_{j,f}}.$$

For a given enzyme loading $f$, the global range-normalized RMSE was computed as

$$\mathrm{NRMSE}_f = 100 \left[ \frac{1}{3|\mathcal{T}|} \sum_{j \in \{\mathrm{Glc}, \mathrm{Xyl}, \mathrm{Man}\}} \sum_{t_i \in \mathcal{T}} \left(\widetilde{\varepsilon}_{j,f}(t_i)\right)^2 \right]^{1/2},$$

where $3|\mathcal{T}|$ is the total number of observations pooled across the three target sugars (Glc, Xyl, and Man) at enzyme loading $f$. This metric quantifies the root mean square prediction error after each residual has been scaled by the observed dynamic range of its corresponding sugar–enzyme-loading data block.

Finally, combined global metrics were computed by pooling the data from both enzyme loadings. The reported global $R^2$ was calculated as the mean of the $R^2$ values for the three target sugars datasets across both enzyme loadings. For the combined range-normalized RMSE, each residual remained normalized by the observed range of its corresponding sugar–enzyme-loading data block:

$$\mathrm{NRMSE} = 100 \left[ \frac{1}{6|\mathcal{T}|} \sum_{f \in \{15,30\}} \sum_{j \in \{\mathrm{Glc}, \mathrm{Xyl}, \mathrm{Man}\}} \sum_{t_i \in \mathcal{T}} \left(\widetilde{\varepsilon}_{j,f}(t_i)\right)^2 \right]^{1/2},$$

where $6|\mathcal{T}|$ is the total number of observations across the three target sugars and two enzyme loadings.

Table S4: Fit-quality metrics for DyAIM, ReAIM, and EAM. Values are reported as NRMSE (%); $R^2$.

| | **15 FPU/g biomass** | | | | **30 FPU/g biomass** | | | | **15 & 30 FPU** |
|---|---|---|---|---|---|---|---|---|---|
| **Model** | **Glc** | **Xyl** | **Man** | **Global** | **Glc** | **Xyl** | **Man** | **Global** | **Global** |
| DyAIM | 1.81; 0.9972 | 2.83; 0.9930 | 3.47; 0.9885 | 2.79; 0.9929 | 2.68; 0.9938 | 3.16; 0.9921 | 2.75; 0.9930 | 2.87; 0.9930 | 2.83; 0.9937 |
| ReAIM | 1.96; 0.9967 | 3.48; 0.9894 | 3.40; 0.9889 | 3.03; 0.9917 | 2.71; 0.9937 | 3.80; 0.9886 | 3.07; 0.9912 | 3.23; 0.9912 | 3.13; 0.9917 |
| EAM | 4.39; 0.9835 | 5.30; 0.9754 | 3.42; 0.9888 | 4.44; 0.9826 | 3.68; 0.9884 | 3.11; 0.9923 | 3.12; 0.9909 | 3.31; 0.9905 | 3.93; 0.9863 |

Table S5: Relative inhibition strengths derived from the optimal DyAIM parameters

| Formula | Description | Value |
|---|---|---|
| $1/K_{I,C}^{G_2}$ | Inhibition potency of $G_2$ on $E_1$ | 0.1679 |
| $1/K_{I,C}^{G_1}$ | Inhibition potency of Glc on $E_1$ | 0.01596 |
| $1/K_{I,C}^{\mathrm{XOS}}$ | Inhibition potency of XOS on $E_1$ | 0.2232 |
| $1/K_{I,C}^{X_1}$ | Inhibition potency of Xyl on $E_1$ | 0.01123 |
| $1/K_{I,C}^{M_1}$ | Inhibition potency of Man on $E_1$ | 0.01073 |
| $1/K_{I,C}^{\mathrm{MOS}}$ | Inhibition potency of MOS on $E_1$ | 0.08749 |
| $1/K_{I,\mathrm{AGX}}^{\mathrm{XOS}}$ | Inhibition potency of XOS on $E_2$ | 0.4682 |
| $1/K_{I,\mathrm{AGX}}^{G_2}$ | Inhibition potency of $G_2$ on $E_2$ | 0.01951 |
| $1/K_{I,\mathrm{AGX}}^{G_1}$ | Inhibition potency of Glc on $E_2$ | 0.00658 |
| $1/K_{I,\mathrm{AGX}}^{\mathrm{MOS}}$ | Inhibition potency of MOS on $E_2$ | 0.09992 |
| $1/K_{I,\mathrm{GGM}}^{M_1}$ | Inhibition potency of Man on $E_3$ during heterogeneous reaction | 1.5022 |
| $1/K_{I,4}^{G_1}$ | Inhibition potency of Glc on $E_4$ | 0.01095 |
| $1/K_{I,4}^{G_2}$ | Inhibition potency of $G_2$ on $E_4$ | 0.01038 |
| $1/K_{I,5}^{X_1}$ | Inhibition potency of Xyl on $E_5$ | 9.9975 |
| $1/K_{I,3}^{M_1}$ | Inhibition potency of Man on $E_3$ during homogeneous reaction | 0.3587 |

### S8 Implementation of the models and the parameter estimation

All three models (DyAIM, ReAIM, EAM) are coded in Python and solved with `solve_ivp` method from `scipy.integrate` module using the BDF integrator (implicit, variable-order 1–5 based on a backward differentiation formula). Parameter estimation, also implemented in Python, uses `least_squares` method from `scipy.optimize` module with the trust-region reflective (`trf`) algorithm. At each stage of the multi-stage optimization procedure, the optimizer minimizes the squared Euclidean norm of the stage-specific residual vector subject to the prescribed parameter bounds, and convergence is assessed with the routine's default gradient and step-size tolerances.

#### S9 Parametric sensitivity analysis

The dynamical models introduced in the present work (DyAIM, ReAIM, EAM) can be written in a state-space form as:

$$\begin{cases} \frac{d\xi}{dt} &= f(t, \xi, \boldsymbol{\theta}), \quad \xi|_{t=0} = \xi_0 \\ y &= g(t, \xi, \boldsymbol{\theta}) \end{cases}$$

where the function $f$ represents the model structure, $\xi = \xi(t, \boldsymbol{\theta})$ denotes the state-space vector and $\boldsymbol{\theta}$ the vector of parameters. The vector of target variables $y = y(t, \boldsymbol{\theta})$ is deduced from the state-space vector thanks to the output function $g$.

The efficiency of the parameter estimation depends on how much the parameters to be estimated influence the target variables. The quantification of this influence is possible thanks to the sensitivity functions defined as [6]:

$$S(t, \boldsymbol{\theta}) = \frac{\partial y}{\partial \boldsymbol{\theta}}(t, \boldsymbol{\theta}).$$

Several methods can be used to compute the sensitivity functions. In particular, sensitivities can be computed by internal differentiation, i.e., by solving the dynamical system provided by the differentiation of the original model with respect to the parameter vector $\boldsymbol{\theta}$ [4, 5]. However, for complex models, this method might require the use of symbolic calculus software. Another method of practical interest, consists of computing the sensitivity functions with the help of a finite-difference discretization with respect to the parameter vector. This is the method that is used in the present work. More precisely, for a given perturbation $\Delta\boldsymbol{\theta}_j$ of the parameter $\boldsymbol{\theta}_j$, the sensitivity of the target variable $y_i$ with respect to the parameter $\boldsymbol{\theta}_j$ is approximated by [1, 2]:

$$\frac{\partial y_i}{\partial \boldsymbol{\theta}_j}(t, \boldsymbol{\theta}) \approx \frac{y_i(t, \boldsymbol{\theta} + \Delta\boldsymbol{\theta}_j e_j) - y_i(t, \boldsymbol{\theta})}{\boldsymbol{\Delta\theta}_j}. \tag{Eq.1}$$

In the above equation, $e_j$ is a vector with the same size as the parameter vector $\boldsymbol{\theta}_j$ such that: all the components of $e_j$ are equal to 0, except the component number $j$ which is equal to 1.

In order to allow comparisons, the sensitivity functions are normalized as follows [4]:

$$(S_{norm})_{i,j}(t, \boldsymbol{\theta}) = \frac{\boldsymbol{\theta}_j}{y_{i,ref}(\boldsymbol{\theta})} \times \frac{\partial y_i}{\partial \boldsymbol{\theta}_j}(t, \boldsymbol{\theta})$$

where $\boldsymbol{\theta}_j$ is the value of the parameter under study and the reference value $y_{i,ref}(\boldsymbol{\theta})$ is chosen as:

$$y_{i,ref}(\boldsymbol{\theta}) = \max_t |y_i(t, \boldsymbol{\theta})|$$

The maximum in the previous equation is taken over all the simulation time points.

In our numerical simulations, the parameter step $\Delta\boldsymbol{\theta}_j$ was chosen as $10^{-4}$ % of the value of the parameter $\boldsymbol{\theta}_j$. The approximation (Eq.1) is achieved by calling twice the Python `solve_ivp` method from `scipy.integrate`. Due to the small value of $\Delta\boldsymbol{\theta}_j$, the high-order solver `rk45` was required in the `solve_ivp` method to avoid numerical instabilities.
